# Transcytosis via the late endocytic pathway as a cell morphogenetic mechanism

**DOI:** 10.1101/2020.01.16.909200

**Authors:** R Mathew, LD Rios-Barrera, P Machado, Y Schwab, M Leptin

**Affiliations:** Directors’ Research Unit, European Molecular Biology Laboratory, 69117 Heidelberg, Germany; Electron Microscopy Core Facility, European Molecular Biology Laboratory, 69117 Heidelberg, Germany; Institute of Genetics, University of Cologne, 50674 Cologne, Germany; National Institute for Science, Education and Research, Bhubaneshwar, India

## Abstract

Plasma membranes fulfil many physiological functions. In polarized cells, different membrane compartments take on specialized roles, each being allocated correct amounts of membrane. The Drosophila tracheal system, an established tubulogenesis model, contains branched terminal cells with subcellular tubes formed by apical plasma membrane invagination. We show that apical endocytosis and late endosome-mediated trafficking determine the membrane allocation to the apical and basal membrane domains. Basal plasma membrane growth stops if endocytosis is blocked, whereas the apical membrane grows excessively. Plasma membrane is initially delivered apically, and then continuously endocytosed, together with apical and basal cargo. We describe an organelle carrying markers of late endosomes and multivesicular bodies (MVBs) that is abolished by inhibiting endocytosis, and which we suggest acts as transit station for membrane destined to be redistributed both apically and basally. This is based on the observation that disrupting MVB formation prevents growth of both compartments.

## Introduction

Most cells have specialized plasma membrane domains that serve dedicated physiological purposes. For instance, epithelial cells have an apical and a basal domain separated by adherens junctions and facing different parts of the body. Membrane and proteins are allocated to these domains in a way that is commensurate with their functions. For example, absorptive epithelia have massively enlarged apical domains organized in microvilli, and photoreceptor cells form specialized membranous outer segments for the light-sensing rhodopsins. Errors in the proportions of membrane domains can have harmful consequences for organ function (Wilson, 2011; Wodarz et al., 1995; Zang et al., 2015). Therefore, the mechanisms that balance plasma membrane distribution are crucial for morphogenesis and tissue homeostasis.

Lipids are synthesized in the ER and trafficked to the plasma membrane via the Golgi apparatus or directly through ER-plasma membrane contact sites (Holthuis and Menon, 2014). Membrane delivery depends on fusion machinery, including SNAREs, and tethers such as the exocyst complex (Saheki and De Camilli, 2017; Wu and Guo, 2015). Due to technical limitations of direct labelling of membrane lipids in vivo, most studies addressing membrane trafficking follow cargo proteins. These are sorted generally at the trans-Golgi network using receptors such as components of the adaptor protein 1 (AP-1) complex (Guo et al., 2014). Rab proteins can also participate in directing polarized secretion (Bellec et al., 2018; Lerner et al., 2013).

Material can also be passed from one domain to the other by transcytosis, which can occur either from apical to basal or vice versa, e.g. IgG and IgA in the gut (Fung et al., 2018). The main role described for transcytosis is to transport cargo from one side of an epithelium to the other. However, redistribution of plasma membrane may also be used for other purposes, including cell morphogenesis (Pelissier et al., 2003; Soulavie et al., 2018). The trafficking routes and the delivery mechanisms are not understood for these processes, nor is it known whether they are isolated special cases.

A cell type with with sophisticated morphogenesis an pronounced specialization of membrane domains is the tracheal terminal cell in insects, which transports oxygen through a branched network of subcellular tubes. Tracheal terminal cells form long hollow branches, with the apical compartment forming the luminal membrane of each branch and the basal compartment facing the body’s inner cavity. This architecture is formed by mechanisms shared by other lumen-forming tissues like endothelial cells and Madin-Darby canine kidney (MDCK) cells grown in 3D (Sigurbjornsdottir et al., 2014). These mechanisms have been widely studied in Drosophila larval tracheal cells (Baer et al., 2012; Ghabrial et al., 2011; Jones et al., 2014; Rios-Barrera et al., 2017; Schottenfeld-Roames and Ghabrial, 2013), where they are however usually limited to endpoint phenotypes with only short-term live imaging possibilities.

By contrast, cell morphogenesis and tube formation can easily be observed live in the embryo (Gervais and Casanova, 2010; JayaNandanan et al., 2014; Okenve-Ramos and Llimargas, 2014; Ricolo et al., 2016), where the first terminal cell branch forms. The subcellular tube develops by an invagination of the apical membrane compartment, and membrane is added throughout the length of the invaginating tracheal tube (Gervais and Casanova, 2010). The tube growths in unison with the basal domain of the cell, and as the branch and its tube extend, trafficking markers are seen throughout the cell, often associated with the tube (Gervais and Casanova, 2010; JayaNandanan et al., 2014; Schottenfeld-Roames et al., 2014). Studies in larvae show that failures in endocytosis, exocytosis and secretion can all affect the growth of both membrane domains as judged by reduced cell branching (Jones et al., 2014; Rios-Barrera et al., 2017; Schottenfeld-Roames et al., 2014). However, only the loss of endocytic components has been associated with defects in subcellular tube size and architecture (Schottenfeld-Roames et al., 2014). How these vignettes of knowledge fit into a picture of an integrated membrane delivery mechanism that balances apical and basal delivery is not known. Here we address this by perturbing membrane dynamics and looking at the redistribution of membrane and protein markers during terminal cell development in the embryo.

## Results

### Organization of membrane domains during subcellular tube morphogenesis

Tracheal terminal cells elongate with their subcellular tube and outer membrane compartment growing at the same rate (Fig. 1A-D). Since lipids are produced in the ER, and membrane proteins and secreted components of the tube must pass through the Golgi, these organelles are likely critical for the expansion of the plasma membrane. To study their distribution we used KDEL::RFP as a maker for ER and the Golgi targeting sequence of β-1,4-galactosyltransferase fused to GFP (GalT::GFP) as a trans-Golgi marker, together with CD4::mIFP, a membrane reporter (Yu et al., 2015) that is enriched in the plasma membrane but also seen in other subcellular membrane compartments. Both organelles are present in the cell body, throughout the length of the growing branch and in the growth cone ahead of the subcellular tube (Fig. 1C, E).

**Figure 1.**
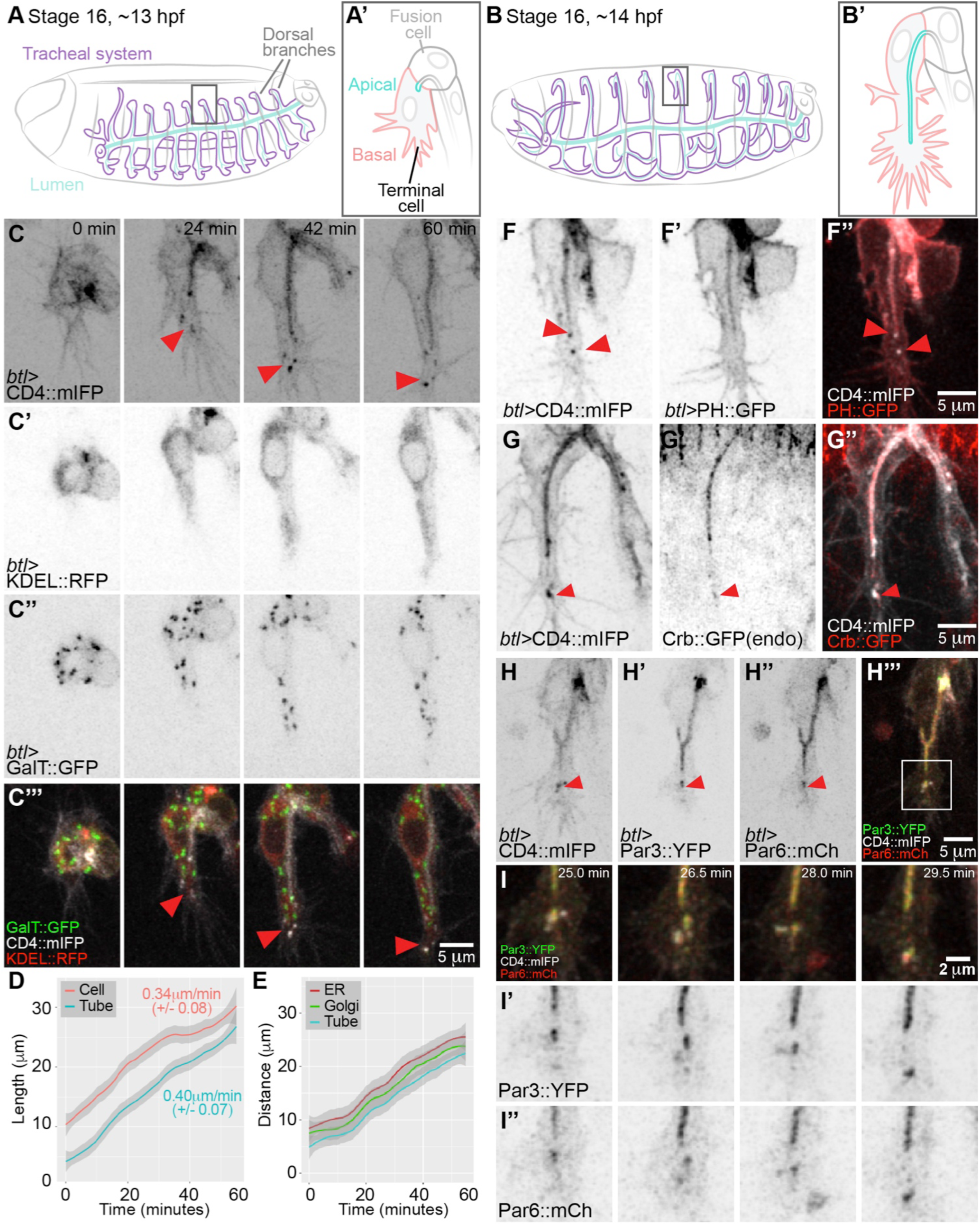
Membrane organization during tube morphogenesis. (A-B) Diagram of the organization of the tracheal system at the onset of terminal cell branching (A) and one hour later (B). Boxed regions are enlarged in (A’-B’). (C-C’’’) Z-projected confocal micrographs of a terminal cell expressing reporters for membrane (C), ER (C’) and Golgi apparatus (C’’). (D) Mean and 95% confidence interval (grey shaded) of tube and branch length during the first hour of growth. Numbers represent the rate of growth estimated from the slope of the curve +/-SD, p=0.2831, *t* test. n=4 (E) Distance of the tip of the tube, and the leading ER and Golgi accumulations from the cell junction that defines the base of the cell, n=6. (F-I) Z-projected confocal micrographs of terminal cells expressing the general membrane marker CD4::mIFP in combination with different membrane or polarity markers: PH::mCherry, a PIP2 sensor commonly used as apical plasma membrane marker (D’), an endogenously GFP-tagged Crb (E’), polarity proteins Par3::YFP (F’) and Par6::mCherry (F’’). The boxed region in (F’’’) is shown at higher magnification in (I) over 4 time points. Red arrowheads show CD4::mIFP vesicles and associated markers at the tip of the cell. In this and the following figures we used the *btl-gal4* line to drive expression of the indicated transgenes. Anterior is left and dorsal is up.

We also noticed accumulations of CD4::mIFP-labelled membrane material in the space between the extending filopodia of the growth cone and the tip of the growing subcellular tube (Fig. 1C). A similar accumulation in this position has been reported for Par3 (which is mainly associated with the apical membrane) but suspected to be an artefact of Par3 over-expression (Gervais and Casanova, 2010). We nevertheless considered the possibility that these structures might be part of the extending tube, or perhaps nascent plasma membrane, and analysed them with a range of membrane markers. They were also seen with other general membrane reporters (Fig. S1A-C), but not with markers considered to be selective for the plasma membrane, such as the Pleckstrin homology domain (PH) of PLCδ fused to GFP or mCherry. These labelled only the tube and the outer, basal membrane of the cell (Fig. 1F, S1B,C, Movie S1). This suggests that the material does not correspond to plasma membrane.

The presence of Par3 in this region is therefore intriguing, and we tested other characteristic apical membrane-associated proteins for their distribution. A GFP inserted into the *crumbs* (*crb*) locus, Crb::GFP, was seen in its normal location at the tube membrane, and also with the CD4 vesicles near the tip of the cell (Fig. 1G), similar to an over-expressed construct (Fig. S1D). Par3 and Par6 also localized to CD4 vesicles ahead of the tube (Fig. 1H). On occasions where different polarity markers were associated with the same vesicle, their overlaps remained partial (Fig. 1I). The fact that Crb::GFP fluoresces indicates that it is not recently synthesized [GFP maturation time is more than 1 hour (Balleza et al., 2018)], and that this compartment therefore does not represent an intermediate along the biosynthetic pathway from the Golgi to the plasma membrane. Instead, we conclude that most likely these structures are endosomes that arise from the apical membrane. This conclusion was supported by high time resolution imaging of Par3 and CD4. The Par3 ahead of the tube appeared to originate from the tube and move to the tip (Fig. S1E, Movie S2), as previously reported (Gervais and Casanova, 2010).

To understand the nature of this domain, we used serial-section electron tomography (Fig. 2, S2). To generate an atlas of organelle distribution throughout the length of the cell we initially screened serial sections manually to identify terminal cells. Though feasible, this labour-intensive workflow allowed only limited analyses. To avoid screening using EM, we turned to correlative light and electron microscopy (CLEM), building on previous protocols that preserve the signal from fluorescent proteins [(Kukulski et al., 2011; Nixon et al., 2009), Fig. S2]. Embryos expressing KDEL::RFP and Par3::YFP under *btl-gal4* were fixed and serially sectioned to cover at least one full embryonic segment (200 sections of 300nm). The fluorescent signal allowed rapid identification of the terminal cells to be imaged by high-resolution electron tomography.

**Figure 2.**
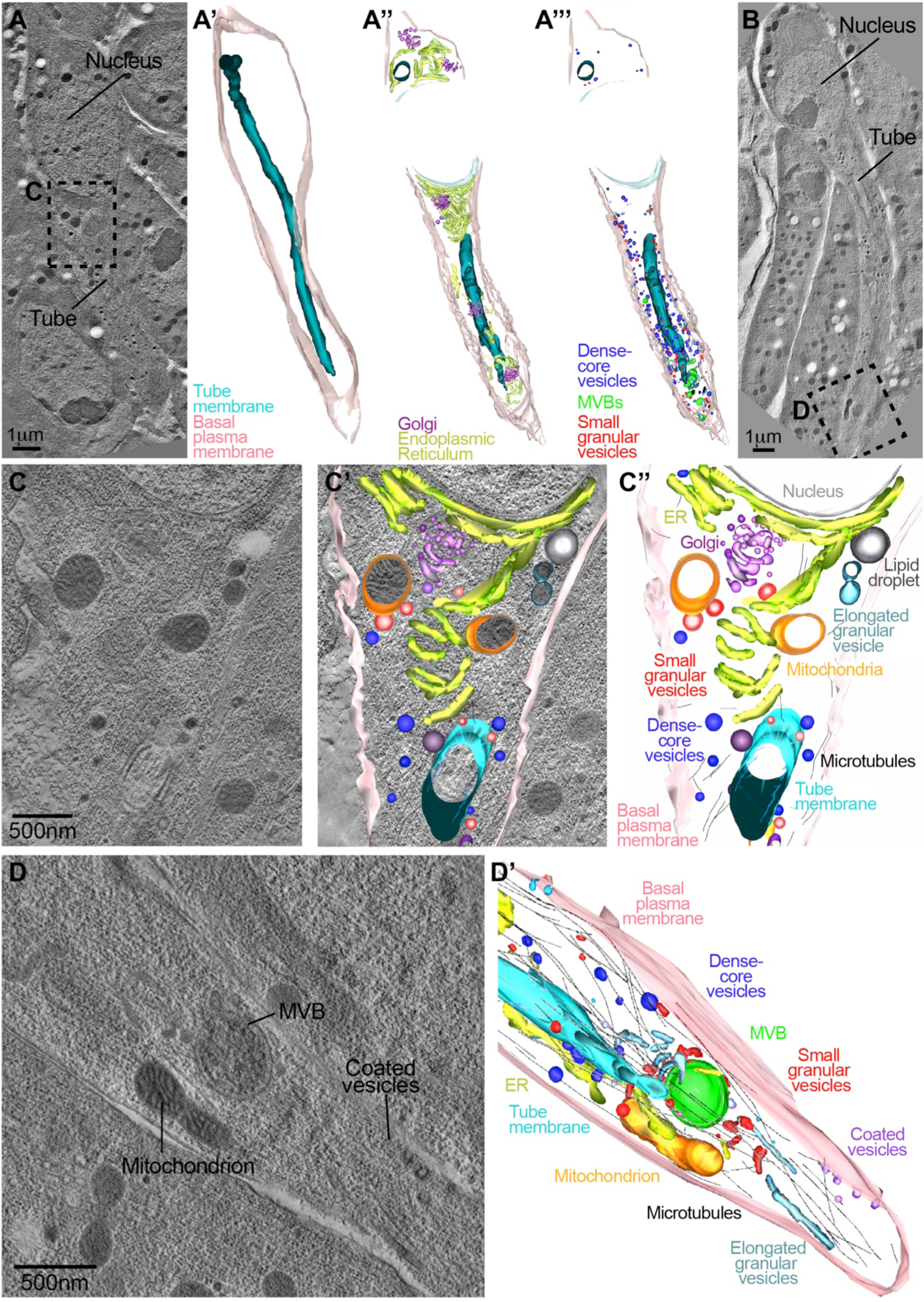
Reconstruction of organelles in a terminal tracheal cell by electron tomography. TEM tomograms from high-pressure frozen embryos. (A-A’) Low magnification (2.5 nm voxels) tomogram and 3D reconstruction from 10 serial 300 nm sections that cover one terminal cell. The dotted box in (A) is shown in (C) at higher resolution. (A’’-A’’’) 3D reconstructions derived from higher resolution tomographs showing the different organelles in the cell. (B) Low resolution tomogram of another cell; the dotted box is magnified in (D). (C-D) Examples of individual high resolution (0.78nm voxels) tomograms used to generate 3D reconstructions. The model in (D’) was derived from 4 serial 3D reconstructions including the one shown in (D).

We first obtained low-resolution overviews of cell morphology (Fig. 2A-B), and then used higher-resolution tomograms to manually trace organelles (Fig. 2C-D, S3A). The tomograms confirmed the distribution of ER and Golgi we had observed by light microscopy, with both organelles spread throughout the length of the cell (Fig. 2A-A’’). We also found a range of vesicles with distinctive electron densities, sizes and distributions (Fig. S3B-C). Many MVBs and large electron-dense vesicles were seen close to the tip of the tube (Fig. 2A’’’,B,D, S3B-C, Movie S3). This, together with the presence of Crb, Par3 and Par6 in this location, suggests extensive membrane trafficking or active recycling events throughout the cell, and particularly in the growing tip.

### The role of endocytosis in terminal cell morphogenesis

The basal plasma membrane and the subcellular tube grow in unison. Hence, there must be a mechanism to balance the delivery of membrane between the two domains. There are in principle two ways to achieve this: 1) Membrane is delivered from the ER and Golgi directly and in the correct measure to each compartment. 2) ER and Golgi deliver membrane to one compartment, and part of this material is then retrieved and transported towards the other via transcytosis. These models can be distinguished experimentally, because the latter requires endocytosis from the plasma membrane for the shuttling. To test this, we blocked endocytosis using a temperature-sensitive allele of *dynamin, shibire*^*ts*^ (Koenig and Ikeda, 1989), which can be inactivated within 15 minutes by shifting the embryos to 34°C.

We blocked dynamin at the onset of tube formation in cells expressing PH::GFP, a construct commonly used as a marker for apical membrane but which is also visible in the basal plasma membrane. Unlike control cells, where basal and apical membrane domains expanded at similar rates (Fig. 3A, C, Movie S4), cells in which dynamin was inactivated failed to grow properly. Measuring fluorescence intensity associated with the outer membrane showed a 1.23-fold increase over 60 minutes (± 0.36 SD, n=5), whereas in the control cells it increased 1.79-fold (± 0.36 SD, p=0.0302). At the same time, *shibire*^*ts*^ cells showed an excessive increase of membrane material inside the cell with a 7.05-fold increase (± 2.98 SD) compared to 2.28-fold of the controls (± 0.5 SD, p=0.0066, Fig. 3B-C, Movie S4). Blocking dynamin function in older cells where the cell and the intracellular tube had already extended led to the accumulation of the marker throughout the length of the subcellular tube (Fig. 3E, Movie S4). The defects in cell and tube growth were reversible: Shifting the embryos back to the permissive temperature restored the expansion of the basal membrane and resulted in partial or complete resolution of the membrane accumulation in the tube domain (Fig. 3B’’, Movie S4).

**Figure 3.**
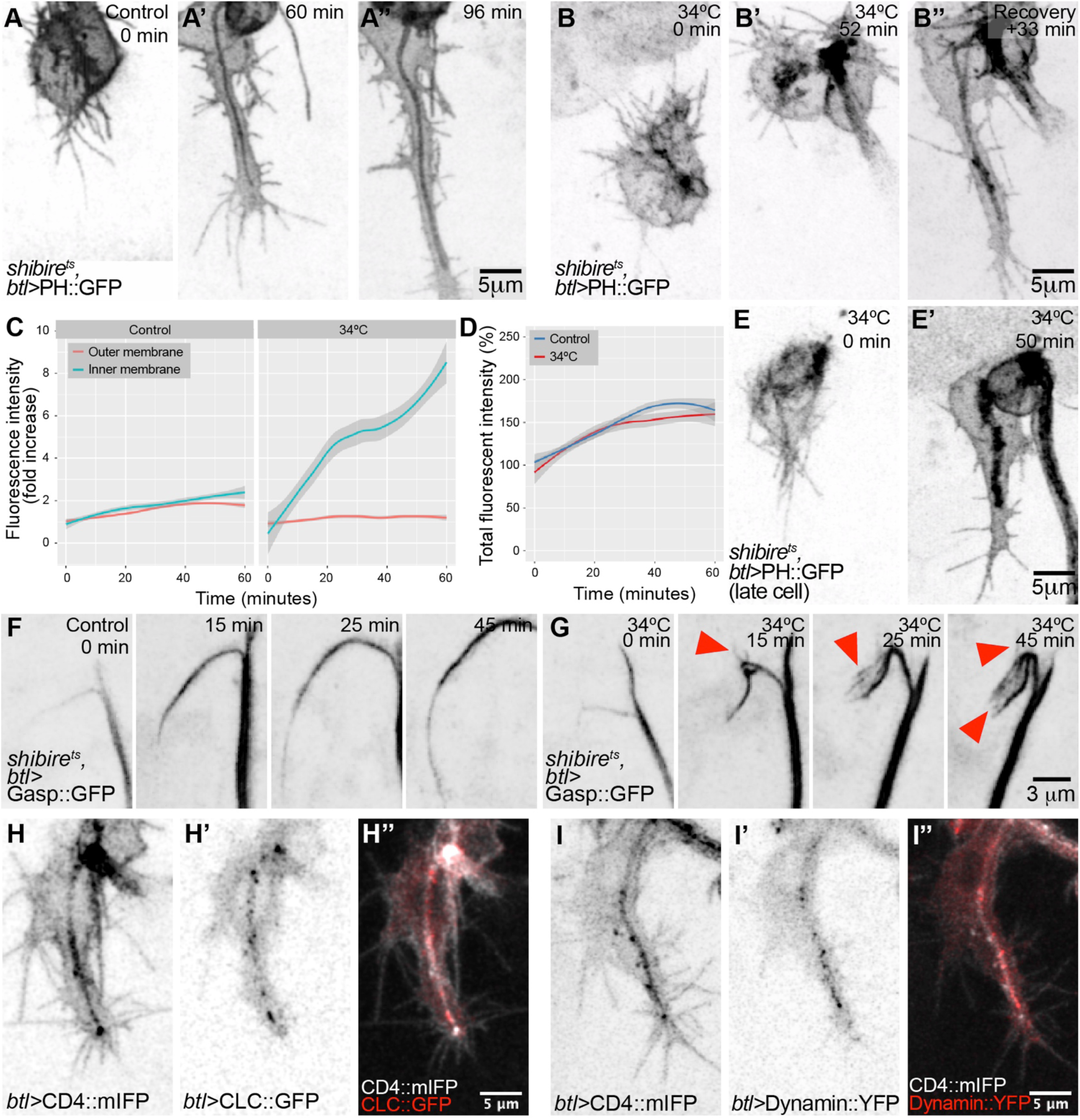
The role of endocytosis in terminal cell growth. (A-E) Distribution of the plasma membrane reporter PH::GFP in control cells (A-A’’) and in cells where dynamin activity had been blocked using a temperature-sensitive allele of *dynamin* (*shi*^*ts*^, B, E). In (B), was inactivated at the onset of tube formation, whereas in (E) it was inactivated ∼15 minutes after the tube had begun to form. (C) Increase in PH::GFP fluorescence intensity in the tube compartment (inner membrane) and in the rest of the cell (outer membrane) over time in control cells (n=7) and also in cells where dynamin was inactivated (n=5). (D) Total fluorescence intensity of PH::GFP increases to the same degree in both conditions. Plots in both panels represent conditional means with 95% CI. (F-G) Distribution of the luminal reporter Gasp::GFP in control cells (F) and in cells where dynamin was inactivated (G). Arrowheads point to protrusions sprouting from the lumen of the terminal cell. (H-I) Distribution of the general plasma membrane marker CD4::mIFP in combination with the Clathrin light chain fused to GFP (CLC::GFP, H) and with Dynamin::YFP (I).

While it was evident that the outer membrane domain expansion was strongly reduced after blocking endocytosis, it was less clear whether the internal membrane pool corresponded to the normal amount that had simply been compacted within a smaller volume, or whether more apical membrane was present. The finding that total fluorescence intensity (basal plus apical) of the cell increased at the normal rate after dynamin inactivation (Fig. 3D) shows that plasma membrane synthesis and delivery *per se* were not affected. Our measurements indicate that upon dynamin inactivation, a similar amount of membrane material as would normally have been added to the basal domain had instead accumulated in the subcellular tube, in addition to the material that is normally delivered there.

We confirmed the identity of the material within the cell as the subcellular tube compartment by the presence of Crb and of Gasp::GFP, a protein secreted into the tracheal lumen. Crb colocalized with the membrane reporter within the cell (Fig. S4B-E), and it remained associated with the tube after recovery (Fig. S4C,F). Gasp::GFP formed complex ramifications that sprout from the subcellular tube (Fig. 3F-G). These results show that dynamin, and therefore most likely endocytosis, is needed for the correct allocation of the appropriate amounts of membrane to the basal and apical compartments. Specifically, membrane is delivered to the apical domain, and it remains there in the absence of endocytosis, while insufficient or no membrane finds its way to the basal domain. According to this model, endocytosis should be more prominent at the apical domain. Consistent with this, both the clathrin light chain (CLC) and dynamin itself were present throughout the cell but they were more abundant around the subcellular tube than in the basal membrane (Fig. 3H-I).

To test whether raised levels of Crb were responsible for the excessive apical membrane, as reported in other contexts (Pellikka et al., 2002; Schottenfeld-Roames et al., 2014; Wodarz et al., 1995), we knocked down Crb (Fig. S4G-H). This did not alleviate the accumulation of apical membrane, and we conclude that blocked endocytosis directly interferes with bulk apical membrane retrieval.

We also considered other potential reasons for the lack of basal membrane growth. Because endocytosis is involved in receptor tyrosine kinase signaling (Villasenor et al., 2016) and FGF signaling is required for terminal cell growth, the growth defect could be due to impaired FGF signaling. To test this, we quantified ERK phosphorylation, a signature of FGF receptor activation (Gabay et al., 1997). Blocking dynamin led to a slight increase in di-phospho-ERK. This was not simply due to the temperature increase, as it was not observed in control embryos shifted to 34°C (Fig. S4I-O). Thus, blocking dynamin function does not prevent FGF signaling activation and if anything, it results in a slight increase in the activation of ERK.

Having excluded other explanations, we suggest that the lack of basal membrane growth and overgrowth of tube membrane are functionally connected, and that the normal balance between the two requires endocytosis. The simplest scenario would be transport of apical membrane material to the basal domain by transcytosis.

### Membrane morphology and distribution in control and *shibire*^*ts*^ cells

To see the effect of blocking dynamin on the membrane or other compartments within the cell we used electron tomography. Control cells had smooth tube membranes, with apical extracellular matrix (aECM) in the lumen of older cells. This appeared as long fibers curling inside the tube and as electron dense depositions adjacent to the plasma membrane (Fig. 4A). In cells fixed after 15 minutes of dynamin inactivation, the tube membrane appeared largely similar to the control (Fig. S5A-B), consistent with the minor effects on cell morphology seen in live observations (Fig. S5C). However, we also found bulges protruding from the tube membrane into the cytosol (Fig. S5B). These were also visible after 1 hour of dynamin inactivation, where additionally the tube membrane appeared highly irregular (Fig. 4B, S5E, Movie S5). We interpret these irregularities as endocytic events in which scission from the membrane had failed to occur, and indeed in several instances these structures were surrounded by particulate electron-dense structures resembling clathrin coats (Fig. 4B’’).

**Figure 4.**
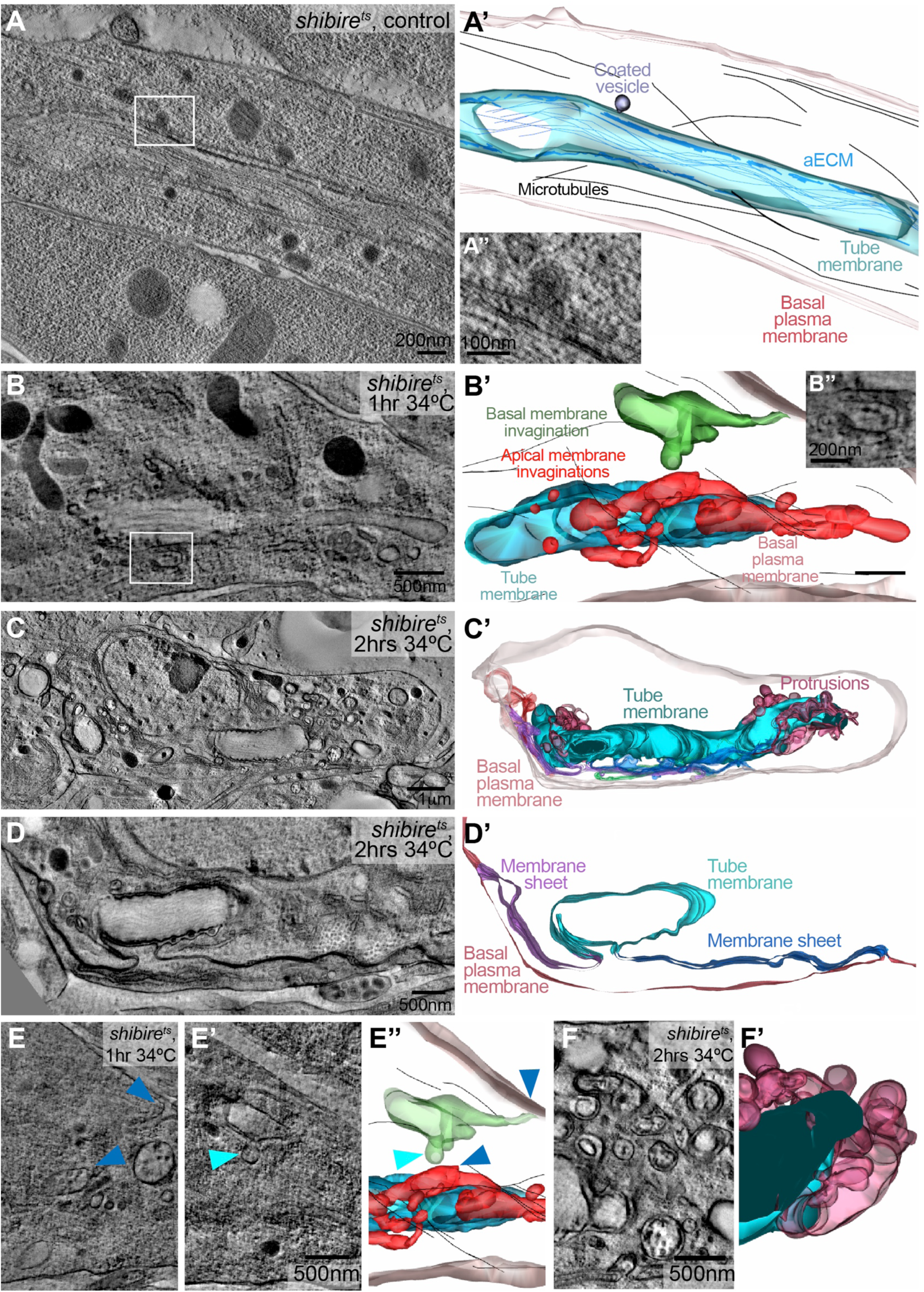
Effects of dynamin inactivation on membrane morphology. TEM tomograms and 3D reconstructions of terminal cells in *shibire*^*ts*^ embryos at permissive (A) and restrictive temperature (B-F). (A-B’’) *shibire*^*ts*^ at permissive (A-A’’) and after 1 hour at restrictive (B-B’’) temperature. Boxed regions in (A) and (B) are magnified in (A’’) and (B’’). They show membrane invaginations surrounded by electron-dense material probably corresponding to a clathrin coat. (A’, B’) Reconstructed 3D models. Color coding of membrane compartments is based on position and structure. The tube membrane (cyan) is characterized by the electron-dense apical extracellular matrix. Invaginations with connections that can be traced to the inner tube membrane are red, invaginations from the outer, basal membrane are green. The basal membrane is light pink. Also shown: microtubules as black lines. (C-D’) *shibire*^*ts*^ terminal cell after 2 hours at restrictive temperature. (C’) 3D reconstruction, color coding for basal and tube membrane is the same as above. In addition, membrane that is continuous with both the apical and the basal membrane is shown in purple and dark blue. (D-D’) higher magnification of a region in the same cell at a level where membrane sheets (purple, dark blue) bridge the apical and basal plasma membrane domains. (E-E’’) Two focal planes and model of a tomogram from the cell shown in (B) where a basal (cyan arrowhead) and an apical invagination are seen in close proximity (closest distance is marked by blue arrowheads). (F-F’) apical membrane overgrowth regions of the cell shown in (C). The genotype of the embryos was *shibire*^*ts*^; *btl>*KDEL::RFP, Par3::YFP. The fluorescent markers were used in the CLEM protocol to identify terminal cells in high-pressure frozen embryos. The cell shown in (C-D) was found and acquired without the CLEM approach.

After 2 hours of dynamin inactivation the morphology of the tube membrane was severely affected. The cells contained complex ramifications of the tube membrane together with its aECM (Fig. 4C, F), which resembled the structures we had seen by light microscopy using Gasp::GFP (Fig. 3G). The cells also showed more dramatic defects. Some invaginations from the tube membrane could be traced all the way to the basal plasma membrane, with which they were clearly contiguous (Fig. 4D, S5F, Movie S5). All cells had instances of large sheets of membrane connecting different parts of the subcellular tube and the plasma membrane. These sheets did not represent autocellular junctions as seen in other tracheal cells (Francis and Ghabrial, 2015) since they contained neither Ecad or FasIII, (Fig. S5G-J). In several places the space between the two apposed plasma membrane sheets contained material resembling the content of MVBs (Fig. 4B, S5F). We view the sheets of membrane bridging the apical and basal membrane as evidence of membrane exchange between the two domains, representing structures that failed to be resolved normally in the absence of dynamin function. By this interpretation, unscissioned membrane invaginations protruding from the subcellular tube would occasionally have touched the basal plasma membrane or its protrusions and fused with it, as transcytosing vesicles would have done in the normal situation. Further plasma membrane delivery may then have expanded such initial channels into larger sheets. This must happen rarely, and when it does happen, rapidly, since we found no clear intermediates in our tomograms, such as apical invaginations reaching to the basal plasma membrane. However, we saw several instances of long invaginations from the apical membrane, occasional ones from the basal membrane, and in one cell, the surfaces of two such invaginations came within 500nm of each other (Fig. 4E, Movie S5).

### Distribution of basal cargo in control and *shibire* cells

If the basal compartment is derived from the apical by endocytosis followed by transcytosis, then general secretion from the ER/Golgi should be directed predominantly towards the apical compartment. This should also apply for transmembrane proteins that are destined for the basal membrane. If secretion follows the route we postulate, blocking dynamin should lead to basal cargo accumulating in the apical compartment before it can reach the basal domain. Hence we looked at the transport of two known basal transmembrane proteins expressed in tracheal cells, the FGF receptor Breathless (FGFR), and the integrin β-chain Myospheroid (βPS-integrin).

In control cells, both proteins mostly localized to the basal filopodia, although FGFR was also seen in large puncta near the tip of the cell (Fig. 5A, S4I, Movie S6). In *shibire*^*ts*^ embryos, both were abnormally accumulated at the tube membrane (Fig. 5B, D, S4O). These findings support our conclusion that the initial delivery of membrane from the Golgi goes mainly to the apical domain, and fails to be redistributed basally in the absence of endocytosis. An alternative interpretation for the apical mislocalisation would be that the defect in *shibire*^*ts*^ cells was not one of failed retrieval of basal proteins, but that cargo was misdirected immediately after the Golgi upon dynamin inactivation, as observed in a mammalian model (Deborde et al., 2008). If this is the explanation for the abnormal localisation of integrin and FGFR, restoring dynamin function should not restore their proper localization. Conversely, if FGFR and βPS-integrin mis-localization was the result of failed membrane endocytosis at the tube, restoring endocytosis should correct their faulty localisation. We observed by live imaging and also by immunostainings that after restoring dynamin function, FGFR distribution to the basal membrane was re-established (Fig. 5B,E, S5M, Movie S6). These results support the model where basal proteins are delivered to the apical membrane first, and are then transcytosed to the basal membrane by dynamin-mediated endocytosis.

**Figure 5.**
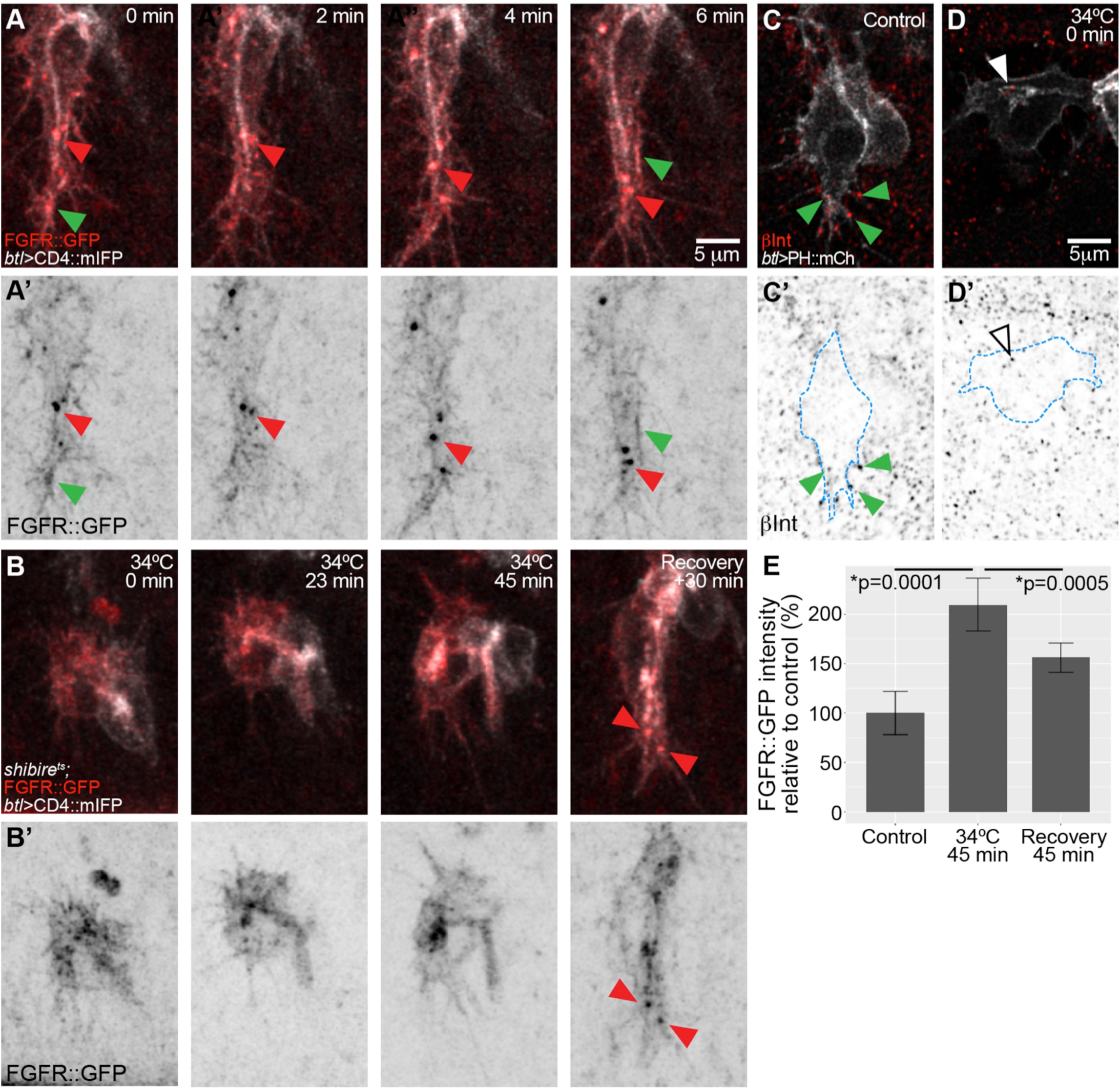
Effect of dynamin inactivation on the distribution of basal proteins. Z-projected confocal micrographs of terminal cells expressing the membrane marker CD4::mIFP under btl-gal4 and FGFR::GFP under its own promoter (from the fTRG library). (A) Time lapse imaging of a control cell. Green arrowheads point to filopodia and basal plasma membrane and red ones point to puncta containing CD4::mIFP and FGFR::GFP. (B) shibirets cell imaged before dynamin inactivation, after 23 and 45 minutes of inactivation, and after 30 minutes of recovery. (C-D) Single confocal planes of terminal cells expressing PH::mCherry and stained for βPS-Integrin. Green arrowheads: βPS-Integrin signal in filopodia; White arrowheads: signal at the tube membrane. The outline of the cell was traced using the PH::mCherry signal and is shown as a blue dashed line in C’ and D. (E) Quantification of FGFR::GFP fluorescence intensity from stained embryos. Number of cells analyzed: Control n=8, Restrictive temperature n=6, Recovery n=8. Significance was assessed using t-test.

### Vesicle trafficking and conversion

Our observations point to a role for apical to basal transcytosis in coupling the extension of apical and basal membrane domains. The ideal way of investigating this experimentally would be to follow the path of plasma membrane lipids by live imaging. Unfortunately no suitable probes exist that allow this in the developing tracheal system *in vivo*. Protein markers as proxies for membranes are not suitable to follow the full path because they are subject to sorting along the vesicle transport path. Instead, we studied individual segments of the potential path by imaging the movement of markers for well-defined vesicular membrane compartments at high temporal resolution. Using CD4::mIFP we found a large number of vesicles moving rapidly along the main axis of the growing terminal cell (Movie S7), often arising in the proximity of the subcellular tube and later moving towards the growing tip of the cell. Vesicle tracking revealed a net distal displacement regardless of where the vesicles originated (Fig. 6A-C). This was not a consequence of cell elongation taking place in the same direction since the vesicles moved faster than the rate of tube elongation (Fig. 6D).

**Figure 6.**
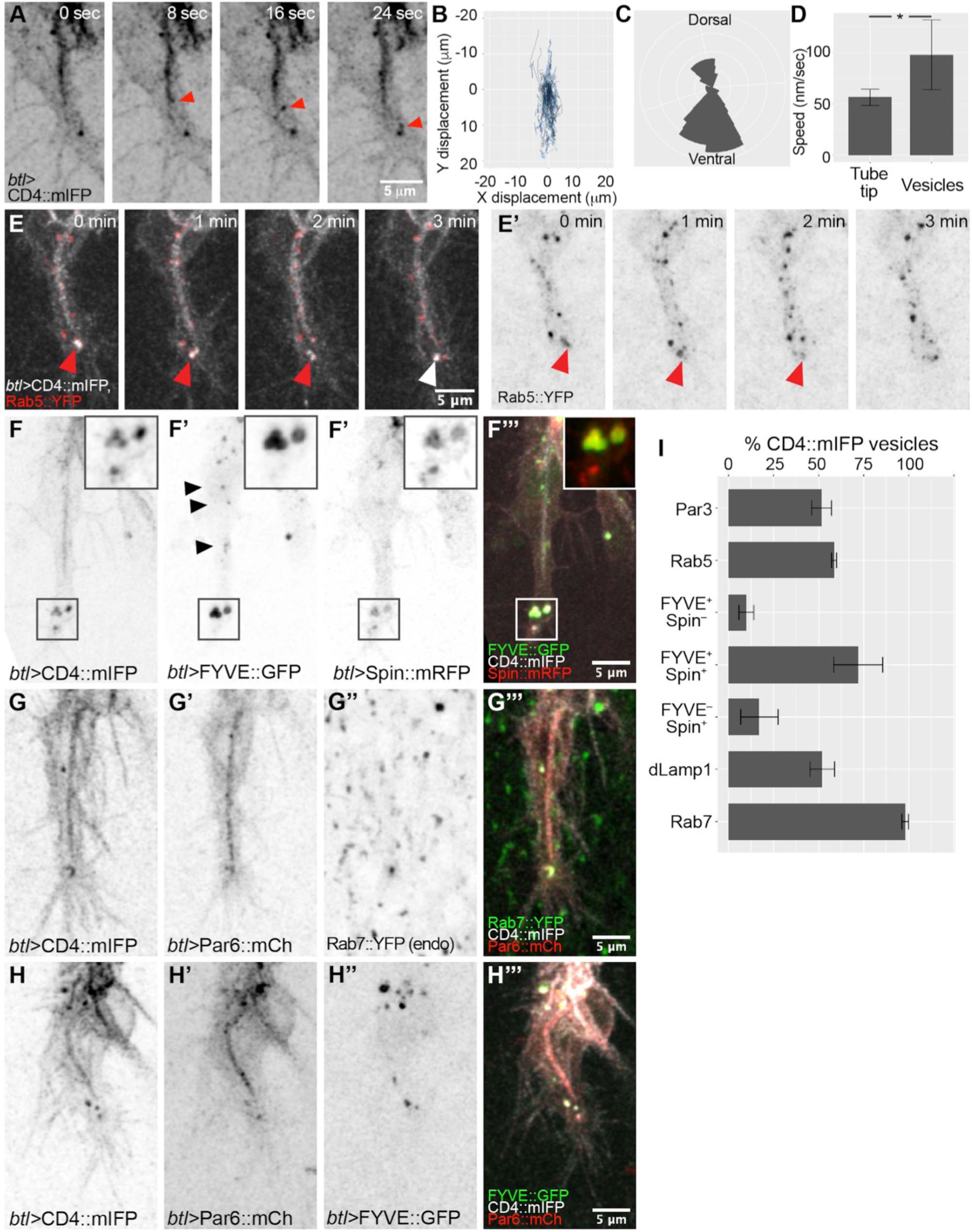
Composition and distribution of endosomal compartments during terminal cell growth. (A-D) High time resolution imaging (8 frames per second) of the membrane marker CD4::mIFP expressed under *btl-gal4*. (A) Example of a large CD4 vesicle (arrowhead) rapidly moving towards the tip of the terminal cell. (B-D) Analysis of 28 CD4 vesicles in five terminal cells. (B) Trajectories of CD4 vesicles with their original position mapped to the origin of the plot. (C) Rose diagram of the trajectories shown in (B). (D) Speed of CD4 vesicles compared to speed of tube growth measured over 25 min; *p<0.0234, *t* test. (E-H). Projected confocal micrographs of terminal cells expressing the membrane reporter CD4::mIFP together with markers for vesicles of the endocytic pathway: (E) Rab5::YFP; (F) the PI3P reporter FYVE::GFP and the lysosomal marker Spin::mRFP; (G) Par6::mCherry and endogenously labelled Rab7::YFP; (H) Par6::mCherry and FYVE::GFP. (E-E’) Red arrowheads: puncta containing CD4::mIFP and Rab5::YFP; white arrowhead: the same CD4::mIFP structure at t = 3min no longer contains Rab5::YFP. (F-F’’’) The boxed regions are magnified in the insets; black arrowheads: small FYVE::GFP vesicles. (I) Proportion of CD4::mIFP vesicles that carry the indicated markers. For each marker we analyzed at least 20 minutes of cell growth. Number of cells analyzed: Par3, n=2; Rab5, n=3; FYVE::GFP, n=4; FYVE::GFP-Spin::RFP, n=3; dLamp1, n=2; Rab7, n=5.

Vesicles first appearing in proximity to the tube suggested that they might be derived from the apical membrane, as was also indicated by the movement of the Par3-positive vesicles (Fig. S1E). We simultaneously imaged CD4::mIFP with GFP fused to the rat atrial natriuretic factor signal peptide (ANF::GFP), an apically-secreted cargo (Tsarouhas et al., 2007). Many CD4 vesicles were positive for ANF::GFP, suggesting that they emerged from the tube (Fig. S6A). In conclusion, we see extensive distal vesicle movement, and at least some of the vesicular membrane appears to be derived from the subcellular tube. To test whether these structures constitute part of the endocytic pathway, we analysed Rab5 distribution as a marker for early endosomes, and found it was visible as discrete puncta around the tube, often localizing to CD4 vesicles. The association between Rab5 and CD4 vesicles was transient, with Rab5 being gradually lost from the vesicles over a few minutes (Fig. 6E). This suggested that Rab5 endosomes were progressing into later endocytic compartments.

Downstream steps of the endocytic pathway involve replacement of Rab5 by Rab7 (Gillooly et al., 2001). Imaging Rab5, Rab7 and CD4 simultaneously showed several instances of CD4 vesicles containing both Rab5 and Rab7, and eventually losing Rab5 while retaining Rab7 (Fig. S6B). Rab5-to-7 conversion results in the recruitment of ESCRT components, leading to MVB formation and afterwards, to the transition into lysosomes. We imaged a range of markers for different steps of the pathway: the FYVE domain of the ESCRT-0 component Hrs (FYVE::GFP) for early endosomes and MVBs, the lysosomal permease Spinster (Spin::RFP), recruited in MVBs, and dLamp1 for lysosomes (Johnson et al., 2015; Riedel et al., 2016). Similar to Rab5, FYVE::GFP formed small puncta throughout the length of the tube (Fig. 6F), but it was more prominently associated with the large CD4-positive structures at the tip of the cell. Spin::RFP, dLamp::mCherry and Rab7 were seen almost exclusively in association with CD4-vesicles at the tip of the cell (Fig. 6F, S6E). Given that the early markers Rab5 and FYVE::GFP were seen throughout the length of the cell, whereas the late markers Rab7, Spin and dLamp were restricted to the tip, we conclude that tube-derived endosomes enter the late endosomal pathway during their movement towards the tip of the cell. This is consistent with our EM analyses where we found a polarized MVB distribution towards the tip of the cell.

To understand the relationship between the tube-derived vesicles, the endocytic pathway and the apical membrane determinants at the tip of the cell, we studied the distribution of apical polarity proteins relative to late endocytic markers. FYVE::GFP, Rab7, and dLamp1 were often seen in proximity to apical proteins (Fig. 6G-H, S6E), suggesting that endocytic vesicles carrying cargo proteins are converted from early to late endosomes as they are displaced towards the tip of the cell.

### Dependence of large intracellular membrane structures on endocytosis

The results so far suggest that the late endosomes at the tip of the cell are sustained by material endocytosed at least in part from the tube membrane. If that was the case, blocking endocytosis should affect them. Quantification of FYVE::GFP-positive vesicles showed that while control cells contained around 1-4 large FYVE::GFP-CD4 vesicles at any given time, cells where dynamin was inactivated had very few, often none, of these vesicles (Fig. 7A-E). Upon recovery, the number of FYVE::GFP-CD4 vesicles was re-established (Fig. 7F-H) and even increased beyond the control conditions. We obtained similar results with Rab7::YFP and Spin::RFP (Fig. S7A-F). This indicates that the large membrane accumulations at the tip of the growing tracheal cell are formed from material delivered by endocytosis. This was consistent with our experiments on Crb and FGFR distribution: both were seen in vesicles in control cells, and these vesicles disappeared upon dynamin inactivation but reappeared during recovery (Fig. S4, 5B).

**Figure 7.**
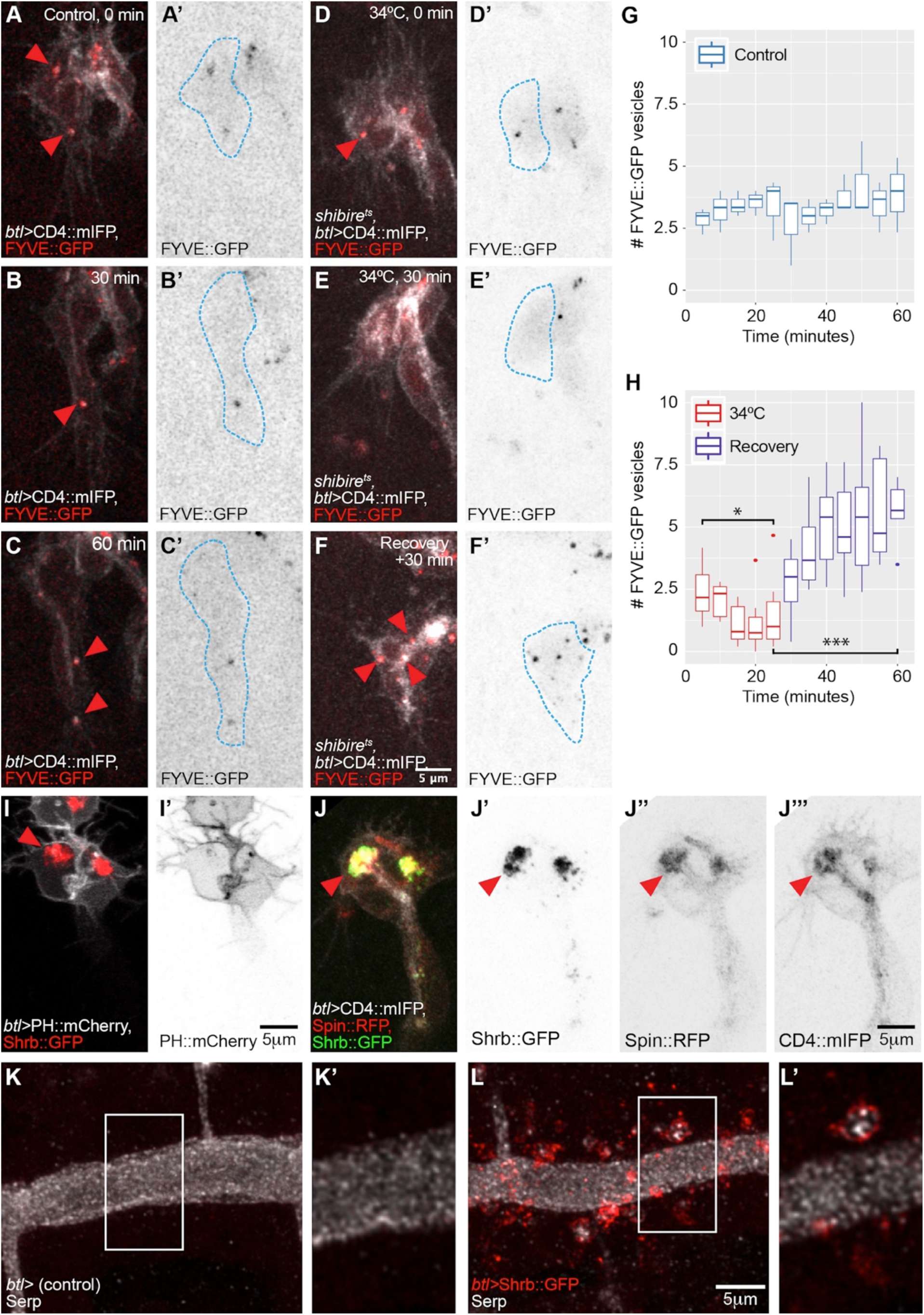
Distribution of late endosomal markers upon disruption of the endocytic pathway. (A-F) Z-projected confocal micrographs of terminal cells expressing CD4::mIFP and FYVE::GFP under *btl-gal4*. (A-C) Control cells. (D-F) *shibire*^*ts*^ cells at permissive temperature (D), after 30 minutes at restrictive temperature (E) and after 30 minutes of recovery (F). Arrowheads: FYVE::GFP puncta. (G-H) Box plots showing the mean number of FYVE::GFP vesicles in 4 control cells (G) and in 6 *shibire*^*ts*^ cells (H). Significance in (H) was assessed using paired *t* test, *p=0.0179, ***p=0.0009. (I-J) Terminal cells over-expressing Shrb::GFP under *btl-gal4* together with PH::mCherry (I), or together with Spin::RFP and CD4::mIFP (J). Arrowheads: Shrb::GFP accumulations. (K-L) Z-projected confocal micrographs of dorsal trunk cells stained for Serp. In control embryos (K), Serp is seen exclusively in the tracheal lumen. The cells themselves are not visible in these images. In embryos expressing Shrb::GFP under *btl-gal4* (L), Serp is also seen inside the cells, usually in association with or surrounded by Shrb::GFP. Boxed regions are magnified in (K’-L’) and shown as single confocal planes.

### Dependence of cell growth on MVB

If the destination of the majority of material endocytosed from the tube membrane is the compartment of late endosomal vesicles at the tip of the cell, the question arises where the membrane material moves from these structures, and whether any of it is delivered to the basal side. MVBs have been shown as a route for delivery of membrane and cargo to the plasma membrane (Dong et al., 2014a; Zhang and Schekman, 2013). Since in our experiments MVBs seem to act as collection points for apical and basal cargo, they may act as a hub for membrane redistribution to the basal membrane domain. To test this, we interfered with the proper biogenesis of the MVBs. MVB formation requires the function of Shrub (Shrb), the Drosophila homolog of ESCRT-III component Snf7, and Shrb::GFP overexpression phenocopies *shrb* loss of function (Dong et al., 2014a; Michelet et al., 2010).

Expression of Shrb::GFP in terminal cells resulted in cells with a morphology similar to that caused by dynamin inactivation: the basal membrane failed to grow, and large accumulations of the CD4 membrane reporter were seen within the cells (Fig. S7G-H). However, unlike in *shibire*^*ts*^ cells these aggregates did not contain PH::mCherry indicating that they were not composed of plasma membrane (Fig. 7I). Instead, the CD4::mIFP-marked membrane accumulation also carried the late endosomal marker Spin::RFP (Fig. 7J). Taken together, this shows that the membrane accumulated inside the cell does not correspond to an overgrown subcellular tube, but rather represents membrane material that was *en route* to MVBs, but because of impaired Shrb function, was not incorporated into MVBs or processed further.

The finding that in this case neither the outer, basal membrane, nor the apical membrane showed any growth indicates that both depend on membrane trafficking through MVBs. For the basal membrane this fits with the observation that its growth depends on endocytosis, but for apical membrane, this was unexpected. It can mean either of two things. All newly synthesized membrane could be directed through the MVB to the apical domain, and the apical domain therefore does not grow if MVBs are disrupted because it never receives membrane. Alternatively, membrane could be passed to the apical domain from the Golgi or the ER directly, but is completely recycled through endocytosis and passage through the MVB. We suggest that only the second scenario is consistent with our observations in *shibire*^*ts*^ cells: The MVBs, as well as other vesicles carrying late endosomal markers disappear in these cells, and yet the tube continues to grow. This shows that the initial delivery of membrane to the apical domain does not require the MVBs. We also confirmed this by blocking endocytosis in cells overexpressing Shrb::GFP, where we found that membrane still accumulated in the plasma membrane even though MVBs were no longer functional (Fig. S7I-J).

These results show that in terminal cells, apical to basal transcytosis goes through MVBs, and we wondered if this was also true for other transcytotic routes. Serpentine (Serp), a chitin-deacetylase located in the tracheal lumen, is produced in the fat body and transported to the tracheal lumen in a processi involving basal to apical transcytosis (Dong et al., 2014b). To test if this process depends on MVBs, we expressed Shrb::GFP in the tracheal system, and looked at Serp distribution in the tracheal dorsal trunk. We found that in control embryos Serp was only seen in the lumen (Fig. 7K-K’). In embryos expressing Shrb::GFP, Serp was also visible in cytoplasmic vesicles that were surrounded by Shrb::GFP itself (Fig. 7L-L’). This suggests that basal to apical transcytosis in the dorsal trunk cells also goes through MVBs.

## Discussion

Our results suggest a route taken by plasma membrane material during the coordinated growth of the apical and basal domains of tracheal terminal cells, summarized in Fig. 8. We will discuss the evidence that leads us to postulate this path, and how it relates to previously discovered functions for components of vesicle trafficking systems and MVBs in cell morphogenesis and transcytosis.

**Figure 8.**
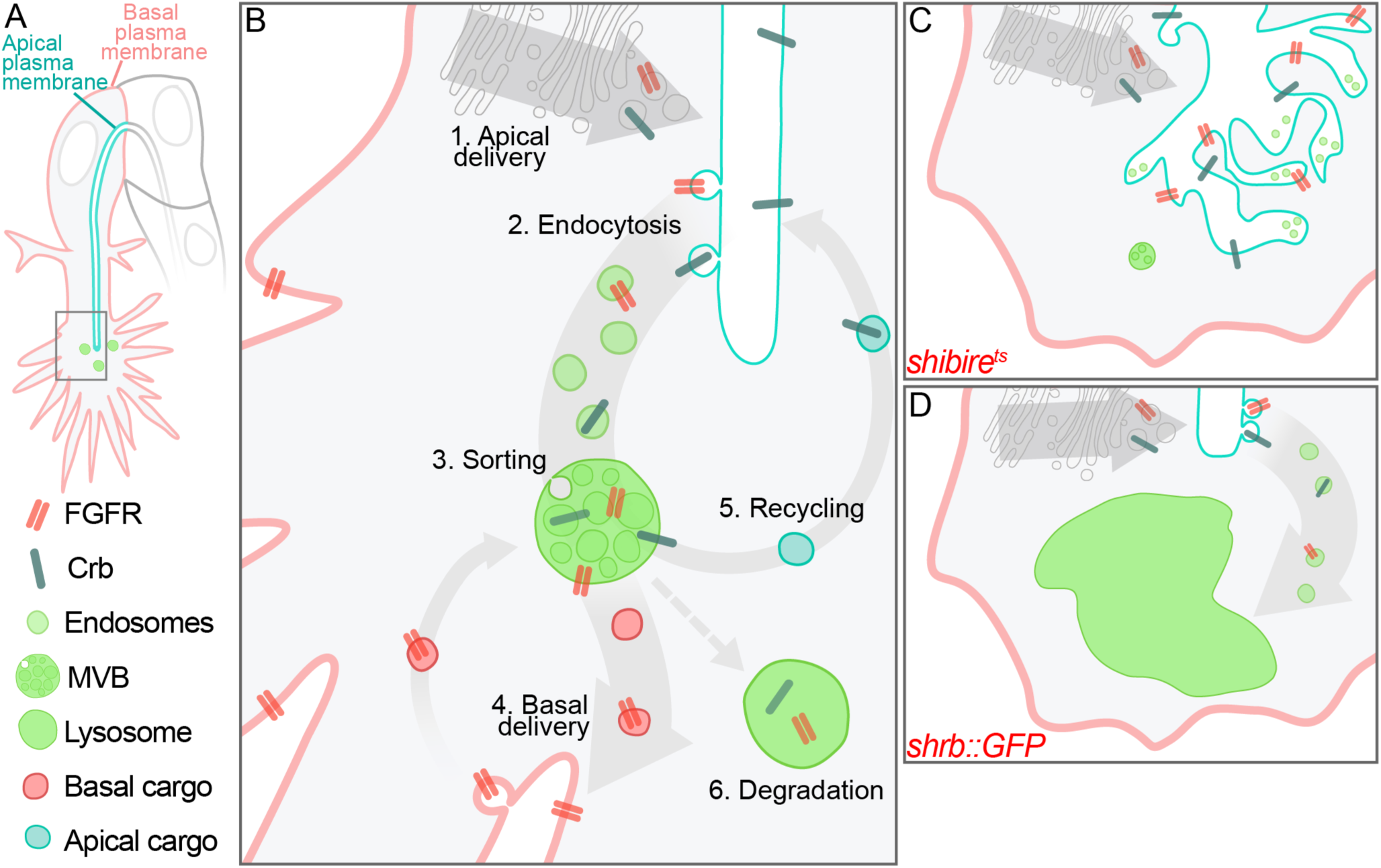
Model of membrane delivery routes during terminal cell growth and tube extension. (A) Overview of a terminal cell and legend of the elements used in (B). The boxed region is magnified in (B). (B) Model for balanced membrane delivery. 1) Membrane and protein material synthesized in the ER/Golgi is delivered apically. 2) Material is endocytosed in a *shibire-*dependent manner, and the resulting vesicles transition from early to late endosomes. 3) As endosomes become MVBs, material is sorted in a *shrb::GFP*-dependent manner. Material is then either transferred to the basal membrane (4), or recycled back towards the apical membrane (5), or targeted to degradation (6). (C) Model of a *shibire*^*ts*^ cell at restrictive temperature. (D) Model of a cell overexpressing Shrb::GFP.

### Initial delivery of membrane

We suggest that apical plasma membrane comes from the ER and Golgi, as also shown in MDCK cells in 3D cultures(Bryant et al., 2010; Ferrari et al., 2008). In terminal tracheal cells, membrane is delivered throughout the length of the tube rather than any specific region (Gervais and Casanova, 2010). This fits with ER and Golgi being distributed along the entire length of the branch. We show that blocking MVB biogenesis prevents membrane delivery to either membrane compartment, which could argue for a role for MVBs in initial membrane delivery. However, we also show that MVB function is downstream of endocytosis.

### Retrieval by endocytosis

Endocytosis plays important roles at multiple steps of tracheal development (Schottenfeld-Roames et al., 2014; Skouloudaki et al., 2019; Tsarouhas et al., 2007), but the fate of the membrane material ingested by the cell through apical endocytosis, the route it takes, and whether it contributes to elaborating the shape of the terminal cell were not known. In larval terminal cells loss or reduction of endocytic components like dynamin, Rab5 or Syntaxin-7 results in cysts in the apical membrane, and sometimes in extreme curling of the apical membrane within branches. These cells also form fewer branches per cell (Schottenfeld-Roames et al., 2014), thus the phenotype suggests an increase in apical membrane at the expense of basal membrane. The presence of excess apical membrane correlated with raised levels of Crb and depended on Crb. In our experiments, even though Crb aggregated at the tube membrane, knocking it down did not prevent apical membrane accumulation. Furthermore, upon recovery of dynamin function, Crb levels were similar to controls. We conclude that here, Crb accumulation does not account for the apical increase resulting from endocytosis being blocked. The difference between the results in the embryo and the larva might be due to extremely different time scales: while we acutely blocked endocytosis for periods of 15 minutes to two hours, larval cells that have grown for several days with reduced levels of endocytic components may be affected by compensatory processes, homeostatic regulation, and longer-term interaction between membrane delivery and Crb accumulation.

### Transcytosis

Transcytosis has been documented in many cell types, most commonly in the transport of proteins and other cargo through epithelial barriers, which can occur either from basal to apical, or vice versa (Callejo et al., 2011; Gallet et al., 2008; Sasaki et al., 2007; Yamazaki et al., 2016). The classical example is immunoglobulin secretion and uptake. In MDCK cells transcytosis of immunoglobulins occurs also in both directions and relies on Rab17 and Rab25. However, similar gene duplications and diversifications have not occurred in Drosophila (Fung et al., 2018; Gramates et al., 2017).

Our conclusion that membrane material is transcytosed from the apical to the basal membrane domain rests on two observations: in the absence of endocytosis the apical membrane continues to grow but the basal stops, and large-scale physical connections appear between the basal and apical membrane after longer periods of blocking endocytosis. These membrane sheets also contain small vesicles similar to the ones produced by the ESCRT pathway to form MVBs, further supporting the delivery route we propose. Consistent with this model, severe dynamin inactivation in larval motor neurons results in a similar phenotype, with membrane cisternae bridging different regions of the synaptic bouton (Kasprowicz et al., 2014).

### Transcytosis as a morphogenetic mechanism

We are aware of only two cases where transcytosis is used as a morphogenetic mechanism: cellularization of the Drosophila blastoderm and morphogenesis of the excretory system in *C. elegan s*(Pelissier et al., 2003; Soulavie et al., 2018). During cellularization, endocytosis retrieves membrane material from the highly folded apical plasma membrane which is subsequently delivered to the leading edge of the growing lateral membranes (Pelissier et al., 2003). The *C. elegans* excretory system forms a subcellular tube similar to the one in Drosophila tracheal terminal cells. Basal to apical transcytosis has been proposed to be required for the growth of this subcellular tube. Tube formation does not depend on dynamin or clathrin but on AFF-1, a protein with no clear homologs in vertebrates or in flies (Sapir et al., 2007; Soulavie et al., 2018). Therefore, it is possible that the excretory canal cell adapted a different strategy to deliver membrane to build its subcellular tube, one that requires AFF-1 instead of clathrin/dynamin-mediated endocytosis.

### MVBs in sorting and membrane delivery

We conclude that trafficking of membrane in the terminal cell goes through MVBs based on our functional results and the colocalization of a number of organelle, membrane, and cargo markers. MVBs are known for their role in protein degradation, especially in the context of ligand-bound receptors (Michelet et al., 2010). However, recycling membrane with its associated transmembrane proteins back to its site of origin forms part of that process and some observations can be interpreted as transcytosing material being associated with MVBs. Gold-labelled IgGs have been traced through various endosomal compartments, including MVBs, before they reach the basal membrane (He et al., 2008). Therefore, it is possible that newly synthesized proteins like FGFR reach MVBs but are then sorted out towards the basal compartment, while others like Crb are translocated into intraluminal vesicles and degraded, if not recycled back to the apical membrane. In agreement with this, in larval tracheal cells loss of ESCRT-0 leads to intracellular accumulation of the FGFR and reduced FGF signal transduction (Chanut-Delalande et al., 2010). According to our model, these results suggest that reduced FGF signaling activity in these cells is due to less FGFR reaching the basal plasma membrane. Thus the defect is upstream of ligand-receptor interactions, rather than in signal transmission after receptor activation.

A multilayered membrane-bound compartment seen in immature larval tracheal branches (Nikolova and Metzstein, 2015) has not been characterized in terms of marker distribution, but it is likely that it is functionally equivalent to the late endosomal compartments that we find at the growing tip of the embryonic terminal cell. Consistent with this model, knock-down of vATPase components in larvae leads to formation of large intracellular Crb vesicles that are Rab5-positive, enlarged Lamp1-positive compartments, and a distorted tube morphology (Francis and Ghabrial, 2015). Cells of the tracheal system that undergo anastomosis rely on specialized late endocytic vesicles called secretory lysosomes to drive tube fusion (Caviglia et al., 2016). These compartments are also positive for Rab7, and their secretion contributes to building a lumen between the fusing cells. MVBs may also be involved in the biogenesis of these vesicles, however downstream mechanisms are likely to differ from the ones employed by terminal cells since fusion cells express a subset of proteins that have no role in terminal cell morphogenesis like Rab39 and the C2 domain protein Staccato (Caviglia et al., 2016).

In summary, our results show that plasma membrane turnover through the late endocytic pathway is a morphogenetic mechanism in which MVBs act as a hub for membrane and cargo sorting. This mechanism entails a massive plasma membrane flow that had so far not been documented. Terminal cells, whose complexity increases dramatically within a few hours in terms of size and morphology, may represent an extreme case of membrane remodeling that relies on this mechanism, but this may nevertheless be required in the building of other complex cell shapes.

## Acknowledgements

We thank Stefano De Renzis, Sandra Iden, Natalia Kononenko, Blanche Schwappach, Catherine Rabouille, and members of the Leptin lab for helpful discussions; the Vienna Drosophila Resource Center, Bloomington Drosophila Stock Center, Kyoto Drosophila Genetic Resource Center and our colleages Markus Affolter, Marko Brankatschk, Stefano De Renzis Chris Doe, Suzanne Eaton, Gabor Juhasz, Elisabeth Knust, Thomas Lecuit, Stefan Luschnig, Christos Samakovlis and Daniel St Johnson for stocks and reagents; the EMBL Advanced Light Microscopy Facility (ALMF) for continuous support, Zeiss for the support of the ALMF, and FlyBase. This work was supported by funding from EMBL and EMBO. LDRB was funded by the EMBL Interdisciplinary Postdoctoral Programme under Marie Curie Actions. We thank Eduardo Rojas-Hortelano for the EM tomogram tracing and analyses.

## Author Contributions

Conceptualization, R.M., L.D.R.B., M.L.; Methodology and investigation, R.M., L.D.R.B., P.M.; Formal Analysis, L.D.R.B; Writing – Original Draft, L.D.R.B.; Writing – Review & Editing, R.M., L.D.R.B., P.M., Y.S., M.L.; Visualization, L.D.R.B.; Supervision, Y.S.; M.L.; Funding Acquisition, M.L.

## Declaration of Interests

The authors declare no competing interests.

## Methods

### Fly lines

*UAS-PLC*δ*-PH-Cherry* was generated by standard molecular biology techniques using UAS-*PLC*δ*-PH-GFP* as template, and cloned into pUAST-attB. The construct was inserted in the VK00033 locus. *btl-gal4* was used to drive UAS transgenes expression in the trachea and was obtained from Markus Affolter lab, University of Basel, Switzerland (3rd chromosome), and Kyoto Drosophila Genetic Resource Center (#109128, 2nd chromosome). The following lines are from Bloomington: *UAS-SrcGFP* (#5432), *UAS-GalT-GFP* (#30902), *UAS-RFP-KDEL* (#30909), *UAS-IVS-myr::tdTomato* (#32221), *UAS-shrbGFP* (#32559), *UAS-GFP-myc-2xFYVE* and *UAS-Spin::mRFP* (#42716), and *UAS-CD4::mIFP-T2A-HO1* (#64182, #64183). From VDRC we obtained *FGFR::GFP* (#318302) and *UAS-crb-IR* (#39177). We are grateful to the groups that kindly shared the following lines: endogenously labelled Rab7 (*YRab7*), and Crb::GFP from Marko Brankatschk, TU-Dresden, Germany (Dunst et al., 2015); *UAS-CLC::GFP, UAS-Shibire::YFP, UAS-Rab5::YFP* and *shibire*^*ts1*^ from Stefano De Renzis, EMBL, Germany (Fabrowski et al., 2013); *UAS-3xeYFP-Baz* (Par3::YFP) from Chris Doe, University of Oregon, USA (Siller et al., 2006); *tub-Rab5::CFP* from Suzanne Eaton, MPI-CBG, Germany (Marois et al., 2006); *dLamp::mCherry* from Gabor Juhasz, Eotvos Lorand University, Hungary (Hegedus et al., 2016); UAS-*crb*^*extraTM*^*::GFP* from Elisabeth Knust, MPI-CBG, Germany (Johnson et al., 2002); *UAS-PLC*δ*-PH::GFP* from Thomas Lecuit, IBDM, France (JayaNandanan et al., 2014); *UAS-palm::NeonGreen* from Stefan Luschnig, University of Münster, Germany (Sauerwald et al., 2017); *UAS-ANF::GFP* and *UAS-GASP::GFP* from Christos Samakovlis, Stockholm University, Sweden (Tiklova et al., 2013; Tsarouhas et al., 2007) and *UAS-Par6::mCherry* from Daniel St Johnston, The Gurdon Institute, UK (Doerflinger et al., 2010).

### Live imaging

Embryos were dechorionated in a 50% bleach solution, washed, and mounted in halocarbon oil using glass coverslips or MatTek glass bottom dishes and heptane glue. Experiments at 34°C were done using a custom incubator chamber mounted on the microscope. For recovery experiments, embryos were mounted in water, incubated at 34°C for the desired time, and for recovery, temperature was lowered to 23°C and ice-cold water was added. Samples were imaged using a Zeiss LSM 880 in Airyscan fast mode, PerkinElmer UltraVIEW VoX or UltraVIEW ERS spinning disc confocal microscopes using Plan-Apochromat 63x/1.4 Oil objectives. Airyscan images were deconvolved using the Zeiss Zen software using the auto settings. Spinning disc images were deconvolved using Huygens Professional, SVI, and processed in FIJI.

### Immunostainings

Embryos were dechorionated, devitellinized and fixed using 37% paraformaldehyde for 15 seconds while vortexing and 5 minutes in a rocker. Afterwards, embryos were blocked using bovine serum albumin and incubated overnight at 4°C in primary antibody solution, and secondary antibodies were incubated for 2 hours the following day. Embryos were mounted in Vectashield. We used the following antibodies: mouse anti-βPS Integrin (1:200, DSHB #6G11), rat anti-ECad (1:100, DSHB #DCAD2), mouse anti-FasIII (1:200 DSHB #7G10), mouse anti-dpERK (1:200, Sigma-Aldrich #M8159), rabbit anti-Dof [1:200 (Vincent et al., 1998)] and rabbit anti-Serp (1:300, gift from Stefan Luschnig, University of Münster, Germany). CBD conjugated to Alexa647 was described previously (JayaNandanan et al., 2014). To enhance signal from fluorescent reporters we used GFP-booster coupled to Atto488 (gba488) and RFP-booster coupled to Atto594 (rba594) from Chromotek. Secondary antibodies used were from Thermo Scientific: Alexa568 goat anti-mouse (1:500, A-11031), Alexa568 goat anti-rat (1:500, A-11077) or Alexa647 goat anti-rabbit (1:300, A-21245). Embryos were imaged using a Zeiss LSM 780 using a Plan-Apochromat 63x/1.4 Oil objective and deconvolved with Huygens Professional and processed in FIJI.

### Electron tomography

#### Initial EM workflow

The embryos were staged and high-pressure frozen (HPM010 AbraFluid) in 20% ficoll (mol weight ∼70,000). The freeze-substitution was done (EM-AFS2 -Leica Microsystems) with 0.3% uranyl acetate, 0.3% glutaraldehyde and 3% water in acetone at −90 °C for 48h. The temperature was then raised to −45 °C at 3.5 °C/h and samples were further incubated for 5h. The samples were rinsed in acetone, followed by infiltration in Lowicryl HM20 resin, while raising the temperature to −25°C. The samples were polymerised under UV light for 48 hours at −25°C and for further 9 hours while the temperature was gradually raised to 20 °C(5 °C/h). Thick sections (300nm) were cut from the polymerized resin block and picked up on carbon and formvar coated slot grids.

The sections were screened using a FEI Tecnai F30 electron microscope with a Gatan OneView camera and acquiring large field of view montages with SerialEM (summarized in Fig. S2). The serial section montages were aligned and segmented using the IMOD package (Mastronarde, D.N. (1997) Dual-axis tomography: an approach with alignment methods that preserve resolution. J. Struct. Biol. 120:343-352.) in order to find terminal cells. The sections that had terminal cells were then imaged again for electron tomography.

#### CLEM workflow

High pressure freezing and freeze-substitution was done as above, except that freeze-substitution was done with 0.1% Uranyl Acetate in acetone at −90 □C for 48h. The fluorescence microscopy imaging of the sections was carried out as previously described (Hampoelz et al. 2016) using a widefield fluorescence microscope (Olympus IX81). The images collected were used to screen for the sections with terminal cells. Those sections were then used for electron tomography. For electron tomography, tilt series were performed in 1 degree increments from 60 to −60 degrees with 2.549 nm or 0,78 nm pixel size on a FEI Tecnai F30 electron microscope with a Gatan OneView camera. The serial tomograms were reconstructed, aligned and segmented using the IMOD package.

#### Image analyses

All images within an experiment were acquired using the same microscope settings. Vesicle tracking and composition, and membrane and tube quantifications were done in FIJI using the manual tracking plugin. For membrane fluorescence intensity calculations we used SUM projections and background subtraction. We then manually segmented the cells and the subcellular tube. For tube fluorescence intensity we subtracted the mean fluorescence intensity of the signal coming from the cell, which should correspond to the basal membrane compartment. For basal membrane fluorescence, we quantified the total fluorescence within the cells and then we subtracted the fluorescence corresponding to the subcellular tube. FGFR and dpERK signal quantifications were done by volume in Imaris, using the Dof signal as mask. Since Dof is a cytoplasmic protein and it does not label fillopodia using it for segmentation allowed us to discard the amount of FGFR still present in the basal plasma membrane.

#### Statistical analyses

We used GraphPad Prism 6 for all statistical analyses. Plots were generated using RStudio and the ggplot2 package.

## References

Baer MM, Palm W, Eaton S, Leptin M, Affolter M. 2012. Microsomal triacylglycerol transfer protein (MTP) is required to expand tracheal lumen in Drosophila in a cell-autonomous manner. J Cell Sci 125:6038–6048. doi:10.1242/jcs.110452

Balleza E, Kim JM, Cluzel P. 2018. Systematic characterization of maturation time of fluorescent proteins in living cells. Nat Methods 15:47–51. doi:10.1038/nmeth.4509

Bellec K, Gicquel I, Le Borgne R. 2018. Stratum recruits Rab8 at Golgi exit sites to regulate the basolateral sorting of Notch and Sanpodo. Development 145. doi:10.1242/dev.163469

Bryant DM, Datta A, Rodriguez-Fraticelli AE, Peranen J, Martin-Belmonte F, Mostov KE. 2010. A molecular network for de novo generation of the apical surface and lumen. Nat Cell Biol 12:1035–1045. doi:10.1038/ncb2106

Callejo A, Bilioni A, Mollica E, Gorfinkiel N, Andres G, Ibanez C, Torroja C, Doglio L, Sierra J, Guerrero I. 2011. Dispatched mediates Hedgehog basolateral release to form the long-range morphogenetic gradient in the Drosophila wing disk epithelium. Proc Natl Acad Sci U S A 108:12591–12598. doi:10.1073/pnas.1106881108

Caviglia S, Brankatschk M, Fischer EJ, Eaton S, Luschnig S. 2016. Staccato/Unc-13-4 controls secretory lysosome-mediated lumen fusion during epithelial tube anastomosis. Nat Cell Biol 18:727–739. doi:10.1038/ncb3374

Chanut-Delalande H, Jung AC, Baer MM, Lin L, Payre F, Affolter M. 2010. The Hrs/Stam complex acts as a positive and negative regulator of RTK signaling during Drosophila development. PLoS One 5:e10245. doi:10.1371/journal.pone.0010245

Deborde S, Perret E, Gravotta D, Deora A, Salvarezza S, Schreiner R, Rodriguez-Boulan E. 2008. Clathrin is a key regulator of basolateral polarity. Nature 452:719–U3. doi:10.1038/nature06828

Dong B, Hannezo E, Hayashi S. 2014a. Balance between apical membrane growth and luminal matrix resistance determines epithelial tubule shape. Cell Rep 7:941–950. doi:10.1016/j.celrep.2014.03.066

Dong B, Miao G, Hayashi S. 2014b. A fat body-derived apical extracellular matrix enzyme is transported to the tracheal lumen and is required for tube morphogenesis in Drosophila. Development 141:4104–4109. doi:10.1242/dev.109975

Ferrari A, Veligodskiy A, Berge U, Lucas MS, Kroschewski R. 2008. ROCK-mediated contractility, tight junctions and channels contribute to the conversion of a preapical patch into apical surface during isochoric lumen initiation. J Cell Sci 121:3649–3663. doi:10.1242/jcs.018648

Francis D, Ghabrial AS. 2015. Compensatory branching morphogenesis of stalk cells in the Drosophila trachea. Development 142:2048–2057. doi:10.1242/dev.119602

Fung KYY, Fairn GD, Lee WL. 2018. Transcellular vesicular transport in epithelial and endothelial cells: Challenges and opportunities. Traffic 19:5–18. doi:10.1111/tra.12533

Gabay L, Seger R, Shilo BZ. 1997. MAP kinase in situ activation atlas during Drosophila embryogenesis. Development 124:3535–3541.

Gallet A, Staccini-Lavenant L, Therond PP. 2008. Cellular trafficking of the glypican Dally-like is required for full-strength Hedgehog signaling and wingless transcytosis. Dev Cell 14:712–725. doi:10.1016/j.devcel.2008.03.001

Gervais L, Casanova J. 2010. In vivo coupling of cell elongation and lumen formation in a single cell. Curr Biol 20:359–366. doi:10.1016/j.cub.2009.12.043

Ghabrial AS, Levi BP, Krasnow MA. 2011. A systematic screen for tube morphogenesis and branching genes in the Drosophila tracheal system. PLoS Genet 7:e1002087. doi:10.1371/journal.pgen.1002087

Gillooly DJ, Simonsen A, Stenmark H. 2001. Cellular functions of phosphatidylinositol 3-phosphate and FYVE domain proteins. Biochem J 355:249–258.

Gramates LS, Marygold SJ, Santos GD, Urbano JM, Antonazzo G, Matthews BB, Rey AJ, Tabone CJ, Crosby MA, Emmert DB, Falls K, Goodman JL, Hu Y, Ponting L, Schroeder AJ, Strelets VB, Thurmond J, Zhou P, the FlyBase C. 2017. FlyBase at 25: looking to the future. Nucleic Acids Res 45:D663–D671. doi:10.1093/nar/gkw1016

Guo Y, Sirkis DW, Schekman R. 2014. Protein Sorting at the trans-Golgi Network. Annu Rev Cell Dev Biol 30:169–206. doi:10.1146/annurev-cellbio-100913-013012

He W, Ladinsky MS, Huey-Tubman KE, Jensen GJ, McIntosh JR, Bjorkman PJ. 2008. FcRn-mediated antibody transport across epithelial cells revealed by electron tomography. Nature 455:542–546. doi:10.1038/nature07255

Holthuis JC, Menon AK. 2014. Lipid landscapes and pipelines in membrane homeostasis. Nature 510:48–57. doi:10.1038/nature13474

JayaNandanan N, Mathew R, Leptin M. 2014. Guidance of subcellular tubulogenesis by actin under the control of a synaptotagmin-like protein and Moesin. Nat Commun 5:3036. doi:10.1038/ncomms4036

Johnson AE, Shu H, Hauswirth AG, Tong A, Davis GW. 2015. VCP-dependent muscle degeneration is linked to defects in a dynamic tubular lysosomal network in vivo. Elife 4. doi:10.7554/eLife.07366

Jones TA, Nikolova LS, Schjelderup A, Metzstein MM. 2014. Exocyst-mediated membrane trafficking is required for branch outgrowth in Drosophila tracheal terminal cells. Dev Biol 390:41–50. doi:10.1016/j.ydbio.2014.02.021

Kasprowicz J, Kuenen S, Swerts J, Miskiewicz K, Verstreken P. 2014. Dynamin photoinactivation blocks Clathrin and alpha-adaptin recruitment and induces bulk membrane retrieval. J Cell Biol 204:1141–1156. doi:10.1083/jcb.201310090

Koenig JH, Ikeda K. 1989. Disappearance and Reformation of Synaptic Vesicle Membrane Upon Transmitter Release Observed under Reversible Blockage of Membrane Retrieval. J Neurosci 9:3844–3860.

Kukulski W, Schorb M, Welsch S, Picco A, Kaksonen M, Briggs JAG. 2011. Correlated fluorescence and 3D electron microscopy with high sensitivity and spatial precision. J Cell Biol. doi:10.1083/jcb.201009037

Lerner DW, McCoy D, Isabella AJ, Mahowald AP, Gerlach GF, Chaudhry TA, Horne-Badovinac S. 2013. A Rab10-dependent mechanism for polarized basement membrane secretion during organ morphogenesis. Dev Cell 24:159–168. doi:10.1016/j.devcel.2012.12.005

Michelet X, Djeddi A, Legouis R. 2010. Developmental and cellular functions of the ESCRT machinery in pluricellular organisms. Biol Cell 102:191–202. doi:10.1042/BC20090145

Nikolova LS, Metzstein MM. 2015. Intracellular lumen formation in Drosophila proceeds via a novel subcellular compartment. Development 142:3964–3973. doi:10.1242/dev.127902

Nixon SJ, Webb RI, Floetenmeyer M, Schieber N, Lo HP, Parton RG. 2009. A single method for cryofixation and correlative light, electron microscopy and tomography of zebrafish embryos. Traffic. doi:10.1111/j.1600-0854.2008.00859.x

Okenve-Ramos P, Llimargas M. 2014. Fascin links Btl/FGFR signalling to the actin cytoskeleton during Drosophila tracheal morphogenesis. Development 141:929–939. doi:10.1242/dev.103218

Pelissier A, Chauvin J-P, Lecuit T. 2003. Trafficking through Rab11 Endosomes Is Required for Cellularization during Drosophila Embryogenesis. Curr Biol 13:1848–1857. doi:10.1016/j.cub.2003.10.023

Pellikka M, Tanentzapf G, Pinto M, Smith C, McGlade CJ, Ready DF, Tepass U. 2002. Crumbs, the Drosophila homologue of human CRB1/RP12, is essential for photoreceptor morphogenesis. Nature 416:143–149. doi:10.1038/nature721

Ricolo D, Deligiannaki M, Casanova J, Araujo SJ. 2016. Centrosome Amplification Increases Single-Cell Branching in Post-mitotic Cells. Curr Biol 26:2805–2813. doi:10.1016/j.cub.2016.08.020

Riedel F, Gillingham AK, Rosa-Ferreira C, Galindo A, Munro S. 2016. An antibody toolkit for the study of membrane traffic in Drosophila melanogaster. Biol Open 5:987–992. doi:10.1242/bio.018937

Rios-Barrera LD, Sigurbjornsdottir S, Baer M, Leptin M, Ríos-Barrera LD, Sigurbjörnsdóttir S, Baer M, Leptin M. 2017. Dual function for Tango1 in secretion of bulky cargo and in ER-Golgi morphology. Proc Natl Acad Sci U S A 114:E10389–E10398. doi:10.1073/pnas.1711408114

Saheki Y, De Camilli P. 2017. Endoplasmic Reticulum-Plasma Membrane Contact Sites. Annu Rev Biochem 86:659–684. doi:10.1146/annurev-biochem-061516-044932

Sapir A, Choi J, Leikina E, Avinoam O, Valansi C, Chernomordik L V, Newman AP, Podbilewicz B. 2007. AFF-1, a FOS-1-regulated fusogen, mediates fusion of the anchor cell in C. elegans. Dev Cell 12:683–698. doi:10.1016/j.devcel.2007.03.003

Sasaki N, Sasamura T, Ishikawa HO, Kanai M, Ueda R, Saigo K, Matsuno K. 2007. Polarized exocytosis and transcytosis of Notch during its apical localization in Drosophila epithelial cells. Genes Cells 12:89–103. doi:10.1111/j.1365-2443.2007.01037.x

Schottenfeld-Roames J, Ghabrial AS. 2013. Osmotic regulation of seamless tube growth. Nat Cell Biol 15:137–139. doi:10.1038/ncb2683

Schottenfeld-Roames J, Rosa JB, Ghabrial AS. 2014. Seamless tube shape is constrained by endocytosis-dependent regulation of active Moesin. Curr Biol 24:1756–1764. doi:10.1016/j.cub.2014.06.029

Sigurbjornsdottir S, Mathew R, Leptin M. 2014. Molecular mechanisms of de novo lumen formation. Nat Rev Mol Cell Biol 15:665–676. doi:10.1038/nrm3871

Skouloudaki K, Papadopoulos DK, Tomancak P, Knust E. 2019. The apical protein Apnoia interacts with Crumbs to regulate tracheal growth and inflation. PLoS Genet 15:e1007852. doi:10.1371/journal.pgen.1007852

Soulavie F, Hall DH, Sundaram M V. 2018. The AFF-1 exoplasmic fusogen is required for endocytic scission and seamless tube elongation. Nat Commun 9:1741. doi:10.1038/s41467-018-04091-1

Tsarouhas V, Senti KA, Jayaram SA, Tiklova K, Hemphala J, Adler J, Samakovlis C. 2007. Sequential pulses of apical epithelial secretion and endocytosis drive airway maturation in Drosophila. Dev Cell 13:214–225. doi:10.1016/j.devcel.2007.06.008

Villasenor R, Kalaidzidis Y, Zerial M. 2016. Signal processing by the endosomal system. Curr Opin Cell Biol 39:53–60. doi:10.1016/j.ceb.2016.02.002

Wilson PD. 2011. Apico-basal polarity in polycystic kidney disease epithelia. Biochim Biophys Acta 1812:1239–1248. doi:10.1016/j.bbadis.2011.05.008

Wodarz A, Hinz U, Engelbert M, Knust E. 1995. Expression of crumbs confers apical character on plasma membrane domains of ectodermal epithelia of drosophila. Cell 82:67–76. doi:10.1016/0092-8674(95)90053-5

Wu B, Guo W. 2015. The Exocyst at a Glance. J Cell Sci 128:2957–2964. doi:10.1242/jcs.156398

Yamazaki Y, Palmer L, Alexandre C, Kakugawa S, Beckett K, Gaugue I, Palmer RH, Vincent JP. 2016. Godzilla-dependent transcytosis promotes Wingless signalling in Drosophila wing imaginal discs. Nat Cell Biol 18:451–457. doi:10.1038/ncb3325

Yu D, Baird MA, Allen JR, Howe ES, Klassen MP, Reade A, Makhijani K, Song Y, Liu S, Murthy Z, Zhang SQ, Weiner OD, Kornberg TB, Jan YN, Davidson MW, Shu X. 2015. A naturally monomeric infrared fluorescent protein for protein labeling in vivo. Nat Methods 12:763–765. doi:10.1038/nmeth.3447

Zang Y, Wan M, Liu M, Ke H, Ma S, Liu LP, Ni JQ, Pastor-Pareja JC. 2015. Plasma membrane overgrowth causes fibrotic collagen accumulation and immune activation in Drosophila adipocytes. Elife 4:e07187. doi:10.7554/eLife.07187

Zhang M, Schekman R. 2013. Cell biology. Unconventional secretion, unconventional solutions. Science (80-) 340:559–561. doi:10.1126/science.1234740

