## Supplemental Figures for "Transcytosis via the late endocytic pathway as a cell morphogenetic mechanism"

**Figure S1. Distribution and composition of membrane reporters during tube morphogenesis.**

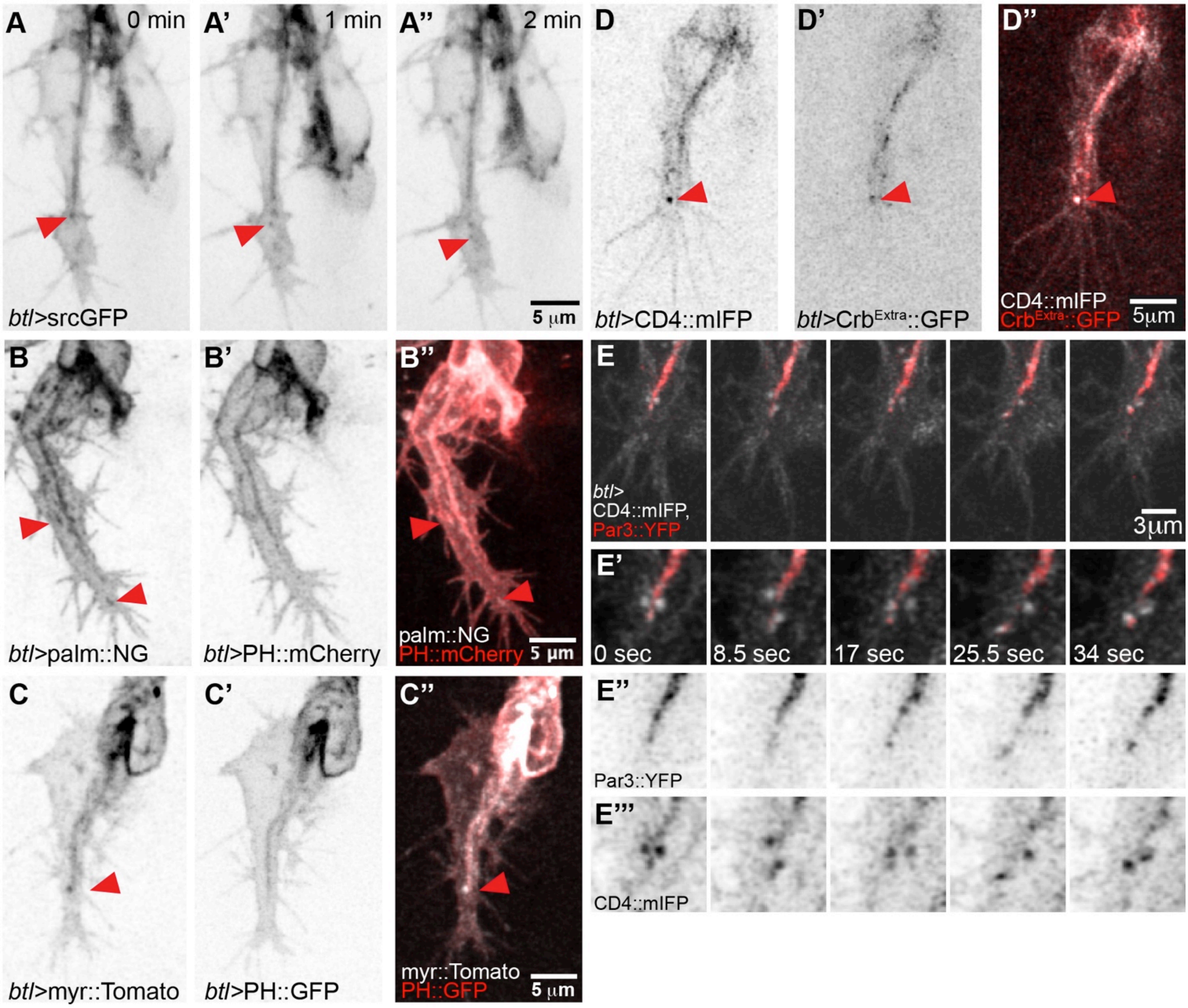


(A-C) Z-projected confocal micrographs of terminal cells expressing membrane reporters: (A-A'') GFP fused to the myristoylation signal of Src, srcGFP; (B-B'') Palmitoylated Neon Green (palm::NG, B') and the PH domain of PLC$\delta$ fused to mCherry (PH::mCherry, B'); (C-C'') myristoylated Tomato (myr::Tomato, C) and the PH domain of PLC$\delta$ fused to GFP (PH::GFP; C'). (D-E) Z-projected confocal micrographs of terminal cells expressing the general plasma membrane marker CD4::mIFP in combination with (D-D'') the extracellular domain of Crb fused to GFP (Crb$^{Extra}$::GFP, D'), and with Par3::YFP imaged at high temporal resolution (E-E'''). (E'-E''') Magnifications of the images shown in (E). Red arrowheads in (A-D) show vesicles and associated markers at the tip of the cell.

**Figure S2. Correlative light and electron microscopy workflow to identify terminal cells**

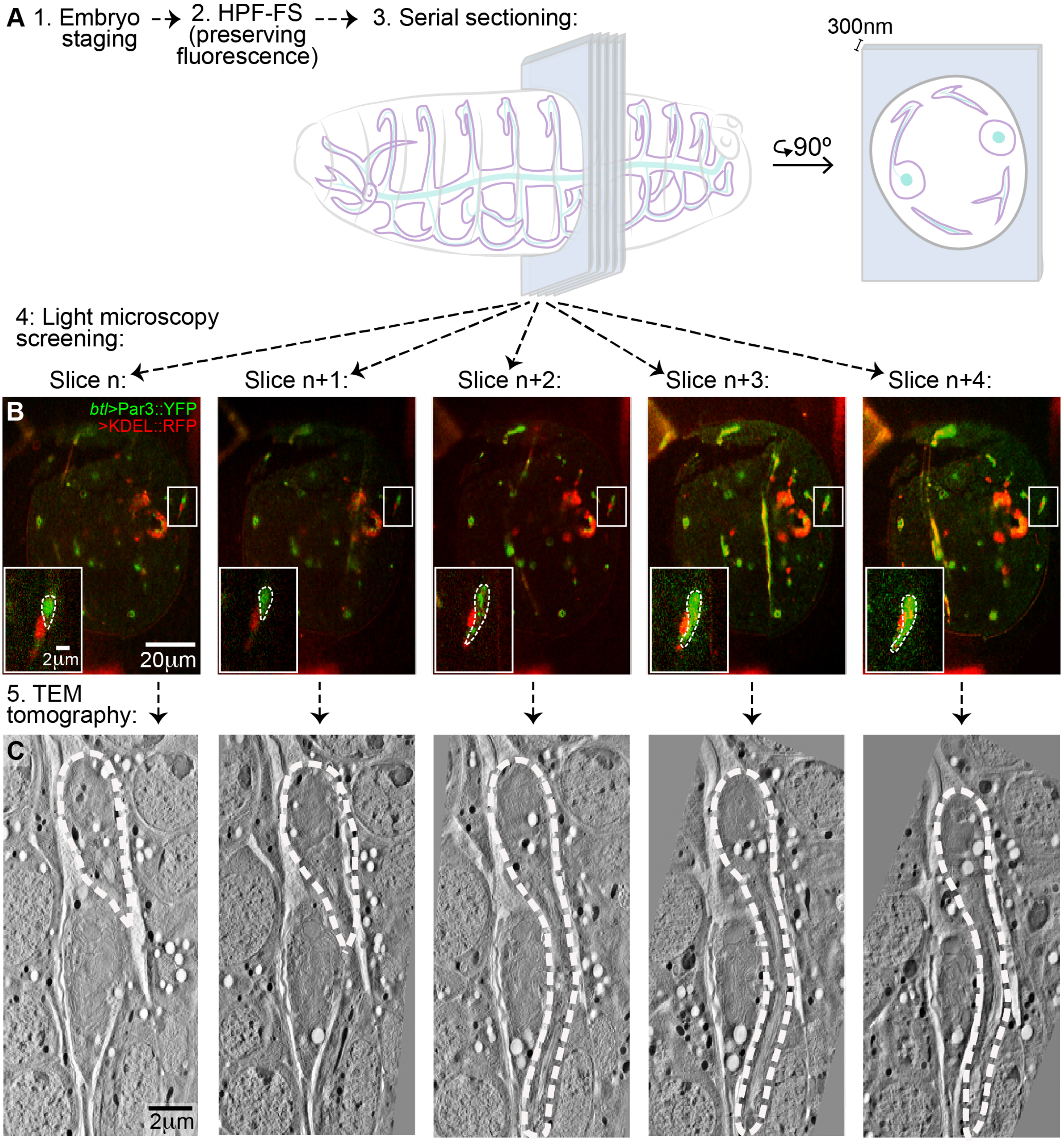


(A) Embryos were processed for EM while preserving the fluorescence signal, and sectioned at 300nm. (B) Physical sections (slices) were then analyzed by fluorescence microscopy and once a terminal cell was identified (Slice n), the adjacent sections were collected to recover the complete cell (Slice n-2 to n+2). (C) The recovered slices were then imaged by electron tomography and digitally aligned.

**Figure S3. Representative vesicle types and their distribution in the terminal cell.**

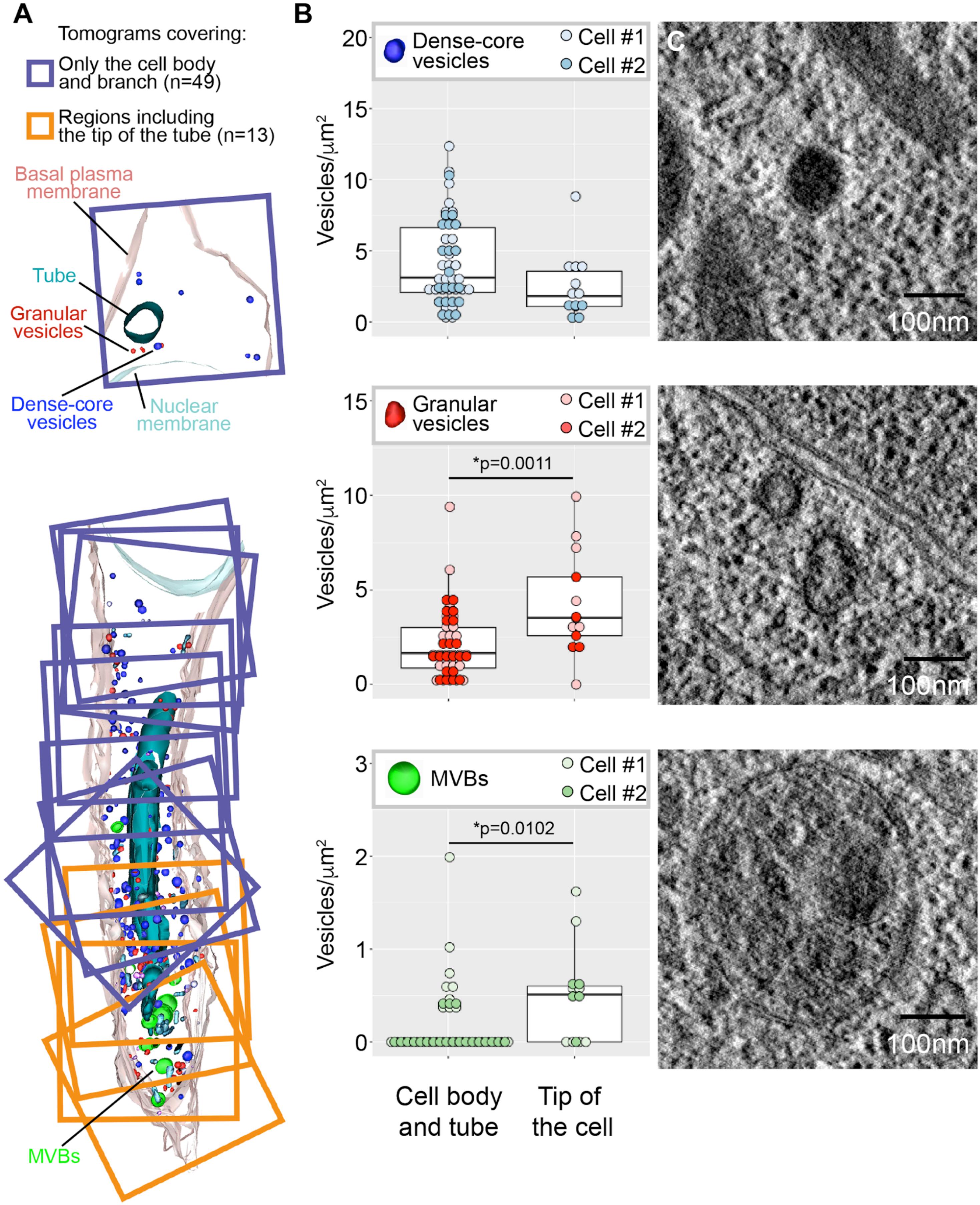


(A) Cell model composed of superimposed 3D reconstructions of high-resolution TEM tomograms covering different regions of the cell. Each square is one 3.2x3.2x0.17 μm 3D reconstruction. The tomograms were assigned to two categories for quantitative evaluation: those including the tip of the tube (orange) and the rest (purple). (B) Distribution of three commonly seen classes of vesicles, expressed as their density in the two regions of the cell (vesicles per $\mu m^2$). Representative examples for each vesicle class are shown next to the quantification plots in (C). Significance was determined using two-tailed *t* test.

**Figure S4. Effect of blocking endocytosis on membrane proteins and FGF signalling.**

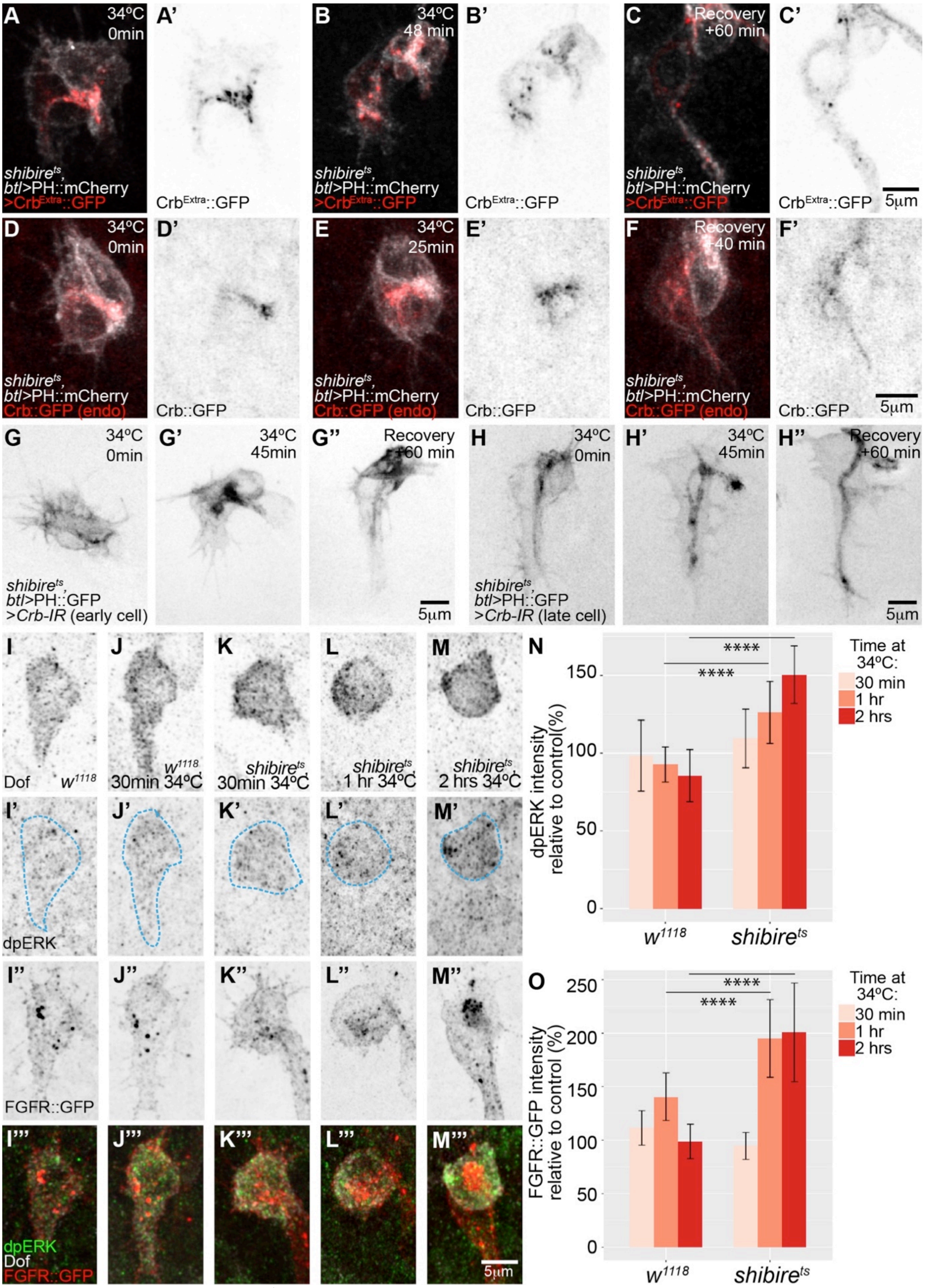


(A-H) Terminal cells of *shibire$^{ts}$* mutant embryos expressing PH::mCherry (A-F) or PH::GFP (G-H). Cells before dynamin inactivation (A, D, G, H), after (B, E, G', H') and at the end of recovery (C, F, G'', H''). (A-C) distribution of the extracellular domain of Crb fused to GFP (Crb$^{Extra}$::GFP); (D-F) distribution of endogenously tagged Crb. (G) Effect of Crb interference RNA (Crb-IR) on a terminal cell at the onset of tube formation. (H) Effect of Crb-IR on a later-stage terminal cell. (I-M) Fixed embryos stained for Dof as a terminal cell marker, dpERK (I'-M', terminal cell position is highlighted in blue), and FGFR::GFP from the fTRG collection (J''-M''). (N-O) Quantification of dpERK and FGFR::GFP fluorescence intensity. Data are plotted as percent relative to control (*w$^{1118}$*, permissive temperature for each time point), +/- SD. We analysed two cells per embryo; number of embryos analysed: *w*, control (for 30 min at 34ºC) n=13; *w*, 30 min at 34ºC n=7; *shi*, control (for 30 min at 34ºC) n=2; *shi*, 30 min at 34ºC n=11; *w*, control (for 1hr at 34ºC) n=18; *w*, 1 hr at 34ºC n=11; *shi*, control (for 1 hr at 34ºC) n=4; *shi*, 1hr at 34ºC n=9, *w*, 2hrs at 34ºCk n=5; *shi* control (for 2hrs at 34ºC) n=6; *shi*, 2 hrs at 34ºC n=6. ****p>0.0001, ANOVA and Tukey's test

**Figure S5. Effects of dynamin inactivation on membrane morphology.**

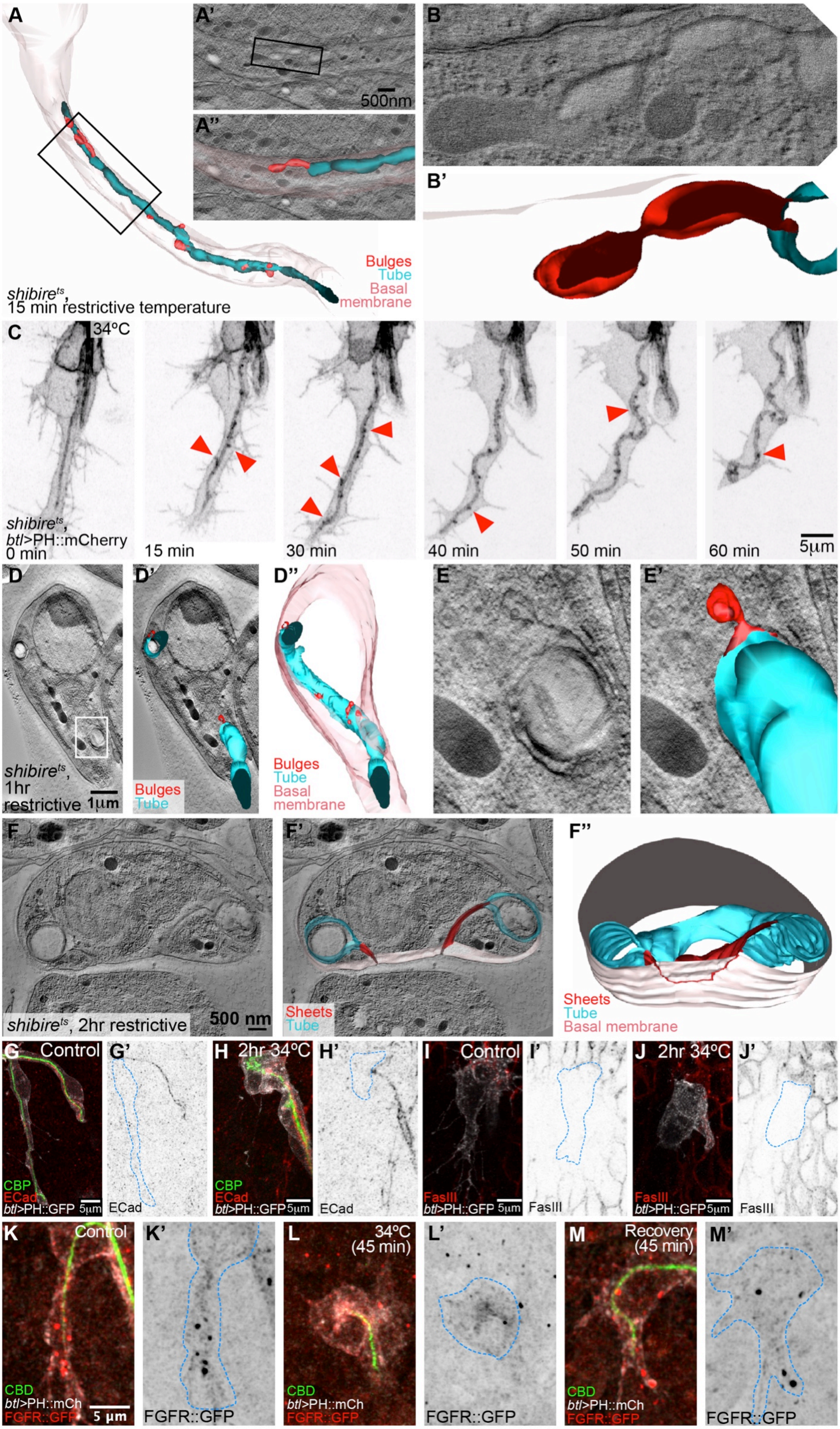

(A, B) Older *shibire*$^{ts}$ terminal cell that had already formed a long branch and tube before dynamin inactivation, and was fixed 15 min after inactivation. (A) Reconstruction of the entire cell. Cyan: apical membrane; red: protrusions from the apical membrane into the cytoplasm, pink: basal membrane. (A', A", B, B') Higher magnification details of the cell, tomograms and reconstructions. (C) Z-projected confocal micrograph of a *shibire*$^{ts}$ terminal cell expressing PH::mCherry. Red arrowheads point to puncta of fluorescent material at the tube membrane. (D-F) TEM tomograms and 3D reconstructions of older *shibire*$^{ts}$ terminal cells similar to (A)?, but after 1 hour (D-E) and two hours (F) at restrictive temperature. Cyan: apical membrane. Pink: basal membrane. Red: protrusions and bulges emanating from the apical membrane, and (F), connecting to the outer, basal membrane. Box in (D) is magnified in (E). (F'') The position at which the sheet between apical and basal is connected to the basal membrane is traced in red on the outside view of the basal membrane. The cells shown in (D-F) were found and acquired without the CLEM approach. (G-J) Z-projected confocal micrographs of fixed *shibire*$^{ts}$ embryos at permissive and restrictive temperature, stained for E-Cadherin and Chitin (CBP, G-H), and for Fasciclin III (I-J). E-Cadherin and Fasciclin are seen at the junctions in the dorsal branch, at the junction of the fusion cell with the terminal cell, but not within the terminal cell. (K-M) Z-projected confocal images of terminal cells stained with a chitin-binding domain fused to a fluorophore (CBD), and stained against PH::mCherry (expressed under *btl-gal4)* and FGFR::GFP (expressed under its own promoter, fTRG library).

**Figure S6. Distribution of late endosomal markers during terminal cell growth.**

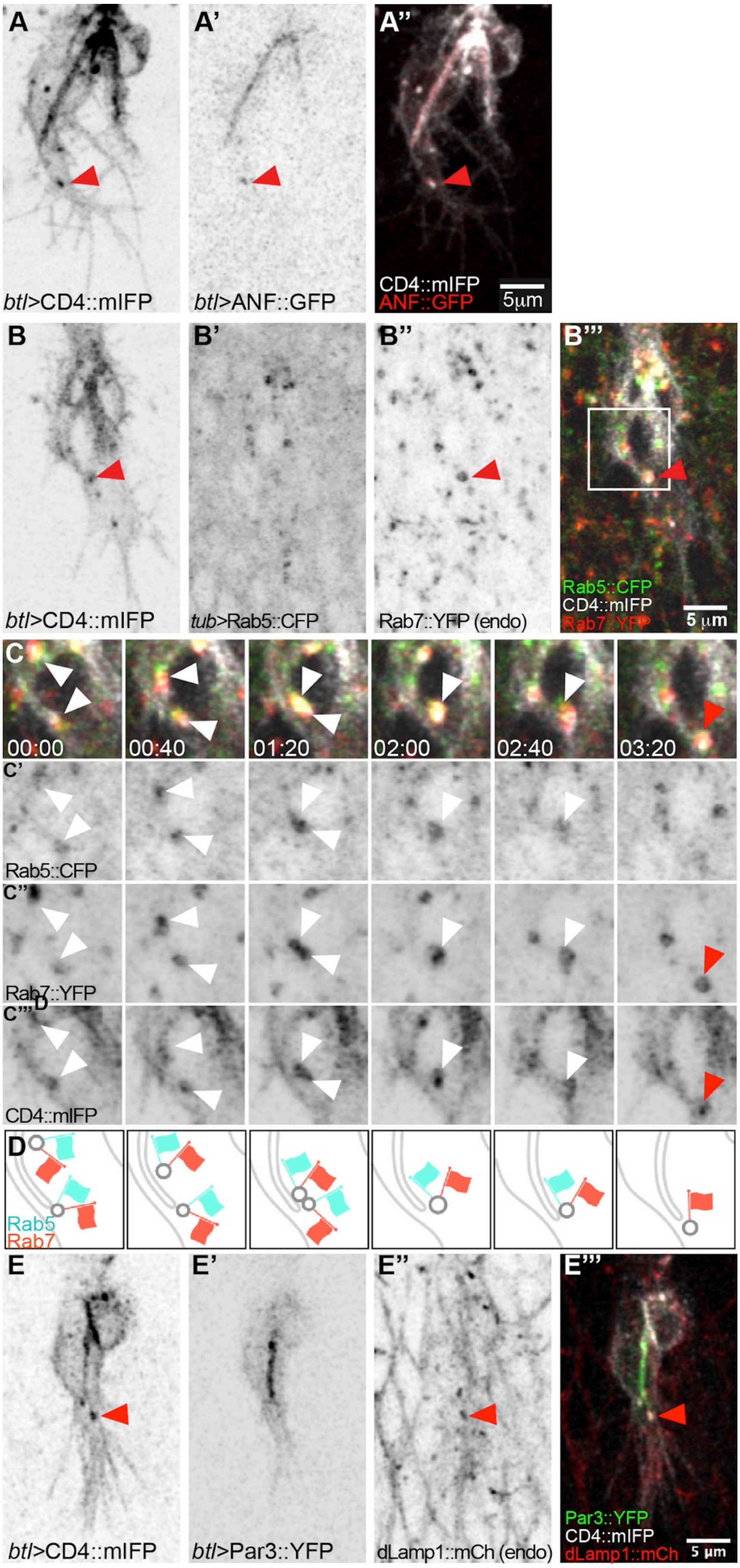


Z-projected confocal micrographs of terminal cells expressing CD4::mIFP under *btl-gal4,* together with late endosomal markers: (A) ANF::GFP; red arrowheads: a CD4::mIFP vesicle that carries ANF::GFP; (B-C) Rab5::CFP under direct control of the tubulin promoter and endogenously tagged Rab7::YFP with Par3::YFP and dLamp1::mCherry under its own promoter. (C) shows the area marked by the box in (B''') at higher magnification and at six time points. (C) White arrowheads: Rab5-positive, Rab7-positive CD4::mIFP vesicles; red arrowhead: Rab5-negative, Rab7-positive CD4::mIFP vesicle. (D) Diagrammatic interpretation of the experiment shown in (C-C'''). (E) dLamp1 presence in a CD4::mIFP vesicle (arrowheads).

**Figure S7. Effect on dynamin inactivation on vesicles carrying late endocytic markers.**

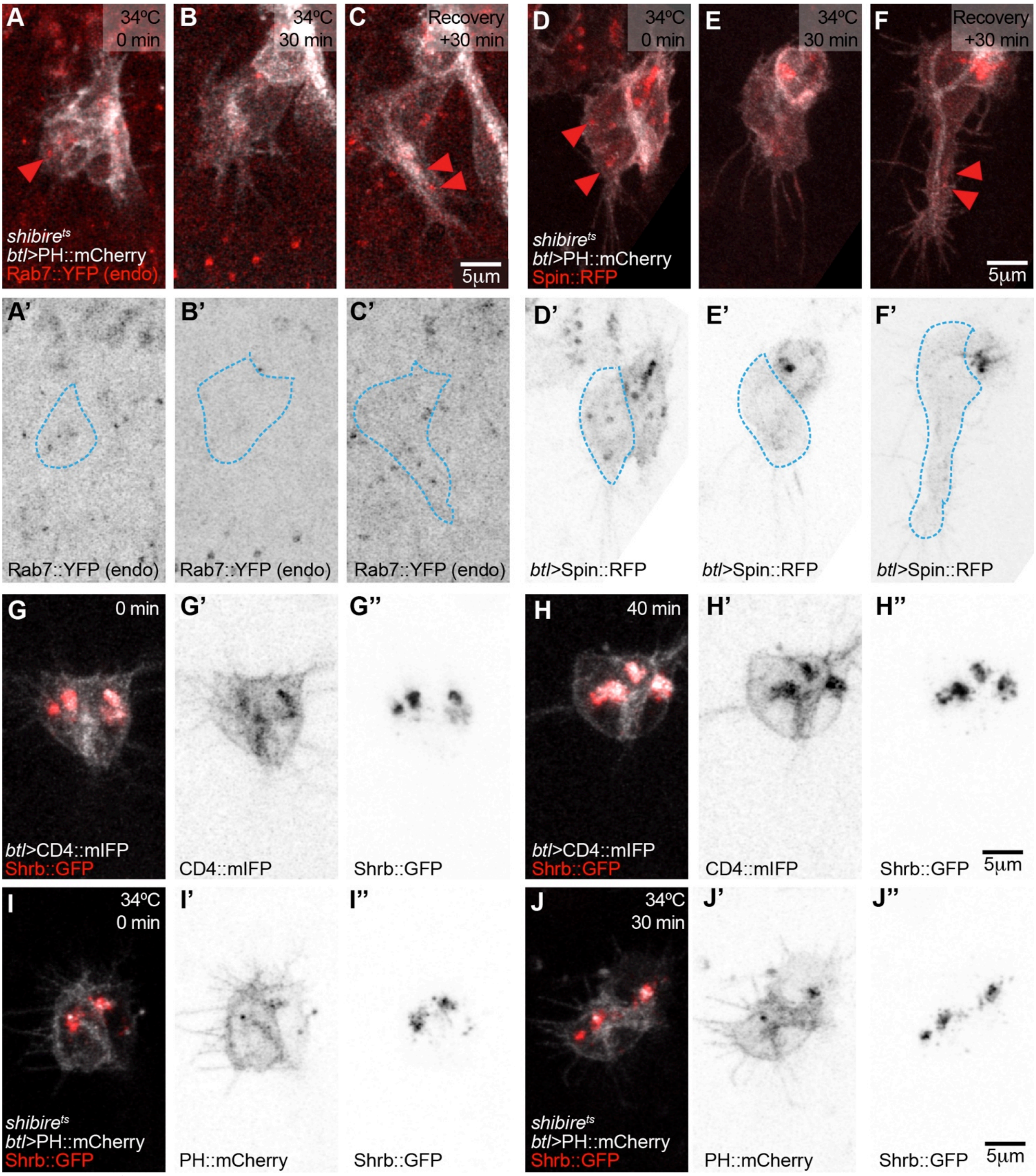


(A-F) *shibire*$^{ts}$ cells expressing PH::mCherry under *btl-gal4*, together with endogenously labelled Rab7::YFP (A-B) or with Spin::RFP (D-F). The outlines of the terminal cells were traced using the mCherry fluorescence and superimposed on the image of the Rab7::YFP channel (blue broken line) to distinguish the cell from the surrounding tissue, which also expresses Rab7::YFP. Arrowheads point to Rab7 and Spin::RFP vesicles in the terminal cell. (G-H) Terminal cell expressing Shrb::GFP and CD4::mIFP under *btl-gal4*, at the onset of tube formation (G) and 40 minutes later (H). (I-J) *shibire*$^{ts}$ terminal cell expressing Shrb::GFP and PH::mCherry under *btl-gal4* before dynamin inactivation (I) and after 30 minutes of inactivation (J).

**Supplemental Video Information**

Movie S1. Distribution of membrane-bound fluorescent reporters
Movie S2. High temporal resolution imaging of *btl*>Par3::YFP and CD4::mIFP
Movie S3. TEM tomograms and 3D reconstruction of a control terminal cell
Movie S4. Effect of dynamin inactivation in terminal cells expressing *btl*>PH::GFP
Movie S5. TEM joined tomograms of terminal cells after 1 and 2 hours of dynamin inactivation
Movie S6. FGFR::GFP distribution in control cells and in *shibire* cells after dynamin inactivation
Movie S7. High temporal resolution imaging of *btl*>CD4::mIFP
